# Ideal Free Distribution in A Multiple Predator-prey System

**DOI:** 10.64898/2026.01.13.699250

**Authors:** Robert Stephen Cantrell, Chris Cosner, King-Yeung Lam, Hua Zhang

## Abstract

We study the mechanism and effects of ideal free distributions (IFDs) on an ecological community consisting of ***n*** prey and ***m*** predator species, for any positive integers ***n*** and ***m***, by considering the corresponding diffusive Lotka-Volterra system with time-periodic coefficients. We define a notion of joint IFD in a time-periodic environment, and give necessary and sufficient conditions for it to be achieved by a subcollection of prey and predator species with suitable dispersal strategies. Next, we show, via construction of a Lyapunov function, that such dispersal strategies are evolutionarily stable, in the sense that if a subcollection of prey and predator species adopts an ideal free dispersal strategy, then the total community must converge to an IFD for large time; if a unique combination of prey-predator species adopts an ideal free strategy, then it can drive all other species to extinction. Conversely, if a combination of prey-predator species adopts a non-ideal free dispersal strategy, then it can be invaded by some suitable mutant strategies. Our results provide insight into the evolution of spatial distribution of ecological communities with predator-prey interactions.

## 1 Introduction

Dispersal is a pivotal life-history trait that fundamentally shapes population dynamics. It mediates interactions between populations, such as predation and competition, and influences the spatiotemporal distribution of individuals. The evolutionary mechanisms underlying dispersal strategies have long fascinated both ecologists and mathematicians, leading to the establishment of several robust theoretical frameworks (Cantrell et al. 2006; DeAngelis et al. 2011; Dockery et al. 1998; Hutson et al. 2001; Lutscher et al. 2005; Křivan and Cressman 2009; Maynard Smith 1974; Geritz et al. 1997; Korobenko and Braverman 2014). In recent years, pairwise invasion analysis, grounded in adaptive dynamics, has become a widely used approach (Averill et al. 2012; Cantrell et al. 2017; He and Ni 2016; Lam and Lou 2014). This method assumes that two (or more) populations differ only in their dispersal strategies, and explore strategies that can resist invasion. The key question is to seek evolutionarily stable strategies (ESS), which are strategies that enable populations adopting these strategies to resist invasion by ecologically identical competitors using any other strategies.

A growing body of research has confirmed that dispersal strategies leading to ideal free distributions (IFDs) are ESS in various settings of dispersal models (Cantrell et al. 2017, 2012, 2022, 2007; Maciel et al. 2020; Cantrell and Cosner 2025; Křivan et al. 2008; Cressman and Křivan 2006). The IFD, due to Fretwell and Lucas (1969), describes how individuals distribute themselves to achieve optimal fitness. This theory is based on two key assumptions: (i) each individual has complete knowledge of its environment, and (ii) each individual can freely move to the most favorable locations. The main motivation behind these assumptions proposed by Fretwell and Lucas was the nesting behavior of birds, which typically have keen eyesight, so they have high abilities to sense the most favorable locations in the surrounding environment (ideal) and then to move to those locations at will (free). Moreover, the IFD prediction has been validated in several empirical or experimental studies (Milinski 1979; Grand 1997; Morris et al. 2004).

The evolutionary impacts of temporal periodicity combined with dispersal in population ecology have garnered significant attention, as most natural systems are influenced by seasonal cycles. Temporal periodicity has been shown to induce significantly different population dynamics. For example, in a two-species competition system with random diffusion, the slower diffuser is favored in temporally constant environments (i.e., slower diffusion is an ESS) (Dockery et al. (1998)). However, in time-periodic environments, the direction of selection can reverse, and coexistence can occur (Hutson et al. 2001; Bai et al. 2023). For further insights into the combined effects of temporal periodicity and dispersal, see Katriel (2022); Liu et al. (2022); Lam and Lou (2024); Benaim et al. (2024); Cantrell and Cosner (2018); Cantrell et al. (2021).

In temporally constant environments, a feature of the IFD is that at equilibrium the reproductive fitness of all individuals will be equal to some constant, which will be zero if the population is at equilibrium. Notably, the occurrence of IFD in temporally varying environments differs from that in temporally constant environments (Kamrujjaman 2019; Lou 2019; Cantrell and Cosner 2018; Cantrell et al. 2021), as they introduce temporal variability of fitness which cannot be avoided no matter what dispersal strategy the population adopts. The simplest example is a single patch population. In this case, if the patch carrying capacity varies with time, then there will inevitably be times when the population overmatches or undermatches the carrying capacity. Consequently, a natural question arises: how to characterize IFD in temporally varying environments. This problem has been considered in a reactiondiffusion-advection model setting by Cantrell et al. (2021) and a patch model setting by Lam and Zhang (2025) with time-periodicity, where an appropriate notion of IFD is introduced based on the concept of pathwise fitness. For general ecological interaction models in time-periodic environments, it is reasonable to conjecture that a similar definition of IFD can be established.

On a broader level, the fitness of a given population depends not only on the local abiotic environment, but also on the collection or the local community of species that it interacts directly or indirectly with (Fauth et al. 1996; McPeek 2017). In fact, the web of interactions among all the species found in a patch of forest, or in a stream or lake, forms very dense and complex networks (Winemiller 1990; Martinez 1991; Dunne et al. 2002; Bascompte et al. 2003). How does the notion of IFD factor into the selection of local community of species? A starting point to address this question is to limit our scope to a smaller subset of interacting species embedded in the broader interaction web, i.e. the community module attributed to Holt (1997).

### The main goals of this paper

In this paper, we continue the study of the problem started in Cantrell and Cosner (2025) concerning the ideal free distribution in predator-prey species interactions in two different ways:

- **(Evolution of a community of species towards IFD)** By introducing a system of equations describing *n*-prey and *m*-predator species, we study how evolution shapes a *community* of prey-predator species by bringing it closer to IFD, where the fitness of a particular prey (resp. predator) species depends on all other members in the community. Roughly speaking, we show that a community of ecologically identical species is stable if and only if it achieves IFD.
- **(IFD in time-periodic environments)** We consider the evolution of IFD in general time-periodic environments. Our argument thus includes as a particular case the evolution of a community to IFD in the autonomous framework as well.

It is worth noting that most current studies on evolution of dispersal primarily focus on two competing species. In such cases, monotone dynamical systems theory can be employed to determine asymptotic dynamics. In contrast, fewer studies have addressed multiple species systems within the framework of adaptive dynamics. Over the course of a lifetime, individuals of any species are likely to interact with multiple competitors, predators, and mutualists. These complex species interactions are crucial for understanding the mechanism of natural selection. Adaptive responses among multiple predator and prey species have been widely recognized as potentially stabilizing forces in large, complex food webs. Consequently, researchers have extensively studied the dynamical behaviors of predator-prey interactions (Flaxman and Lou 2009; Sadovsky and Senashova 2016; Schreiber and Saltzman 2009; Cantrell and Cosner 2025; Cantrell et al. 2007, 2012; Křivan 1996; Cressman et al. 2004; Zelenchuk and Tsybulin 2021; Frølich and Thygesen 2022). The IFD of predator-prey systems has been explored in Cantrell and Cosner (2025); Cantrell et al. (2007, 2012); Křivan (1996); Cressman et al. (2004); Zelenchuk and Tsybulin (2021); Frølich and Thygesen (2022). However, to our knowledge, the majority of the applications of the adaptive dynamics approach to IFD in predator-prey systems are limited to environments that are static in time.

Unlike the existing work, we apply pairwise invasion analysis to investigate the evolutionary stability of IFD of a multiple predator-prey species model in time-periodic environments. The presence of multiple species raises an intriguing question: if each single species adopts a non-ideal free dispersal strategy, but the superposition of their distribution constitutes an ideal free dispersal strategy, would the so-called *joint IFD* strategy be selected or not?

### 1.1 A reaction-diffusion model consisting of *n* prey and *m* predator species

We will study this problem through the following Lotka-Volterra predator-prey model with *n* prey species and *m* predator species:

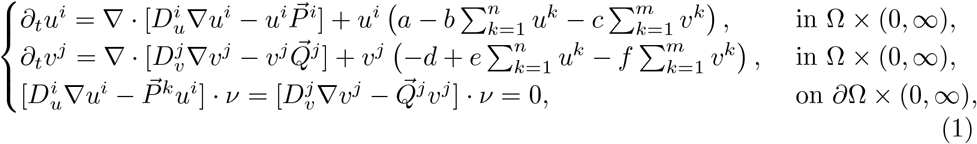

where Ω is a bounded smooth domain in ℝ^*N*^, *i* = 1, …, *n* and *j* = 1, *m*. Here, function *u*^*i*^(*x, t*) (resp. *v*^*j*^(*x*, …*t*)) represents the density of the *i*-th prey species (resp. *j*-th predator species) at time *t >* 0 and location *x* ∈ Ω, where the set Ω is a bounded domain in ℝ^*N*^ with a smooth boundary *∂*Ω and an outward unit normal vector *ν*.

The dispersal strategy of prey species *u*^*i*^ is given by 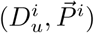, where 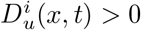 is the spatial diffusion rate and the vector field 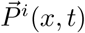 describes the directed movement. A similar interpretation holds for the predator species with dispersal strategy 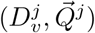. The functions *a*(*x, t*) and *d*(*x, t*) represent the intrinsic growth rate of a prey species and the death rate of a specialist predator species, respectively. The functions *b*(*x, t*) and *f*(*x, t*) measure the strength of self-limitation, and *c*(*x, t*) and *e*(*x, t*) are the predation rate and the production rate of predator species, respectively.

Since we are primarily concerned with the evolution of dispersal strategies, we assume that the collection of prey (resp. predator) species are interacting neutrally, which is indicated by the appearance of ∑_*k*_ *u*^*k*^ (resp. ∑_*k*_ *v*^*k*^) in the demographic rates, and that the coefficients *a, b, c, d, e, f* are independent of *i* and *j*, so that the prey (resp. predator) species differ only by their dispersal strategies.

We recall the following hypotheses in Cantrell and Cosner (2025):

**(Ha)** *f*(*x*) *>* 0 in 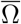 and there exists a constant *κ*_1_ *>* 0 such that

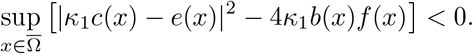

**(Hb)** *f*(*x*) ≡ 0 in 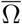 and there exists constant *κ*_2_ *>* 0 such that *e*(*x*) = *κ*_2_*c*(*x*).

**(Hc)** There is a constant *C*_*v*_ ∈ [0, 1) such that

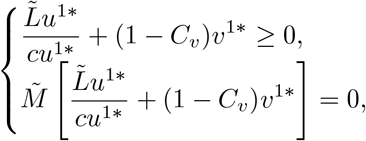

where 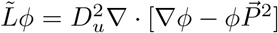 and 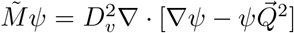 and (*u*^1*^(*x*), *v*^1*^(*x*)) is uniquely determined by

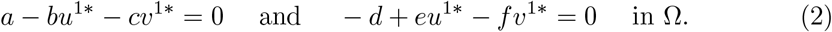

#### Theorem 1

*(Cantrell and Cosner 2025) Consider system* (1) *with time-independent coefficients, where m* = *n* = 2 *and* 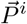 *and* 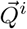 *are the gradients of some functions. Suppose that the first predator-prey pair adopts an ideal free dispersal strategy, while the second pair does not, then the following statements hold*.

a. *If* **(Ha)** *holds, then the steady state u*^1*^(*x*), 0, *v*^1*^(*x*), 0 *given in* (2) *is globally asymptotically stable among nonnegative and nontrivial initial conditions*.
b. *If* **(Hb)** *holds, but* **(Hc)** *does not, then the steady state* (*u*^1*^(*x*), 0, *v*^1*^(*x*), 0) *is globally asymptotically stable among nonnegative and nontrivial initial conditions*.

The second closely related result concerns a model of three competing species written as follows:

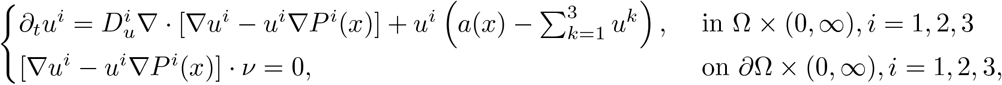

Where 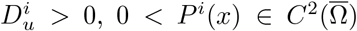, and *a*(*x*) is positive and non-constant. It is easy to see that if exp(*P* ^1^(*x*)) = *k*_0_*a*(*x*) for some *k*_0_ *>* 0, then the corresponding 5 equilibrium of the single species exactly matches the availability of resources, i.e. *u*^*i*^(*x*) ≡ *a*(*x*), and hence is an IFD. Similarly, we say that (*P*^*i*^, *P*^*j*^) forms an IFD pair if *a*(*x*) ∈ span {exp(*P*^*i*^), exp(*P*^*j*^)}. Then it is proved in Theorems 2.2 and 2.3 (Lou and Munther 2012) that

i. If species 1 adopts an ideal free dispersal strategy and species 2 and 3 do not form an ideal free pair, then species 1 excludes species 2 and 3.
ii. If species 1 and 2 form an IFD pair, and species 3 does not adopt an ideal free dispersal strategy, then species 1 and 2 exclude species 3.

Theorem 1 indicates that predator-prey species adopting ideal free dispersal strategies drive those with non-ideal free dispersal strategies to extinction, while the results of Lou and Munther (2012) demonstrate that species adopting ideal free dispersal strategies or collectively adopting such strategies will dominate during multi-species competition. A similar result has been proven in Cantrell et al. (2012), which considers an arbitrary number of competing species in temporally constant, patchy environments.

Inspired by the above results, we inquire what are the selection mechanisms behind the dispersal when including time periodicity. Specifically, we will generalize the definitions of IFD in Cantrell and Cosner (2025); Cantrell et al. (2021) and ideal free pairs in Lou and Munther (2012) to system (1), where multiple interacting prey-predator species in a time-periodic environment are considered, see Definitions 1 and 2. We then perform a pairwise invasion analysis to determine whether a subset of prey species {1, …}, *n* and a subset of predator species {1, …}, *m* using an ideal free dispersal strategy can resist invasion by the remaining species using other dispersal strategies. The time periodicity brings a challenge for constructing Lyapunov functions for (1). We handle this problem with the generalized relative entropy method (Lam and Lou 2022, Chapter 4).

Throughout this paper, we make the following assumption:

(**A**)The coefficients are smooth and *T*-periodic in *t*, i.e.,

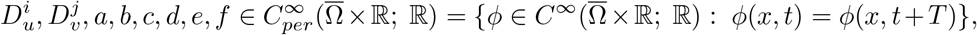

and 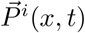, and 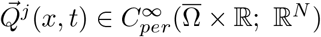, and such that

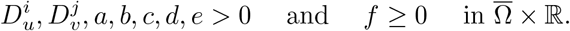

### 1.2 The definition of joint IFD for system (1)

Motivated by Definition 2.1 in Cantrell et al. (2021), we introduce a notion of IFD for a predator-prey system in a time-periodic environment:

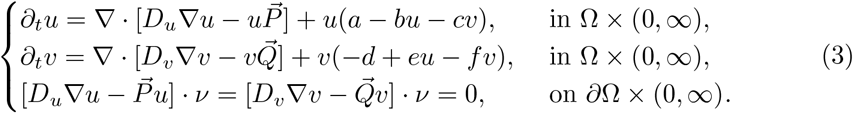

#### Definition 1

For a positive, *T*-periodic solution (*u*^*^, *v*^*^) of system (3), we say (*u*^*^, *v*^*^) is an IFD if

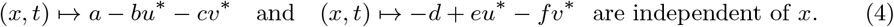

Moreover, we say the corresponding dispersal 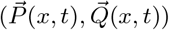 is an ideal free dispersal strategy.

*Remark 1* In case the coefficients are temporally constant, then (4) is consistent with our previous definition of IFD.

*Remark 2* Consider the special case where the coefficient functions satisfy

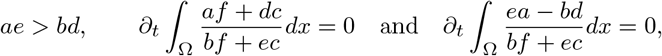

then define, for each *t*, the functions *p*_0_(*·, t*), *q*_0_(*·, t*)

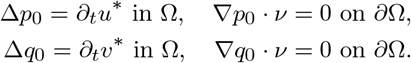

One can verify that the dispersal strategy

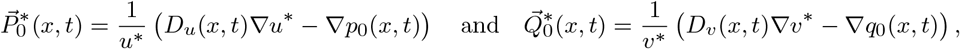

gives rise to the *T*-periodic solution 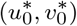 of (3) given by

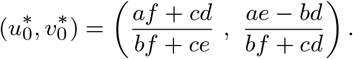

This is an IFD since 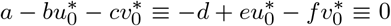, i.e., 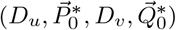 is an ideal free dispersal strategy.

As shown in Lou and Munther (2012); Cantrell et al. (2012), it may happen that a specific combination of distributions for multiple species exactly form an IFD. This leads to the notion of a joint IFD for predator-prey systems with arbitrarily many species, which is a generalization of Definition 1.

#### Definition 2

Let 𝒮 be a nonempty subset of {1, …, *n*} and 𝒦 be a nonempty subset of {1, …, *m*}. We say that system (1) has a 𝒮𝒦 joint IFD 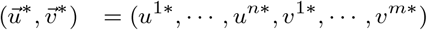 if 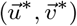 is a *T*-periodic solution of (1) satisfying

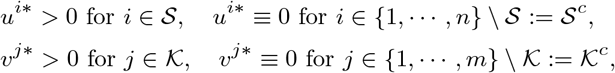

and

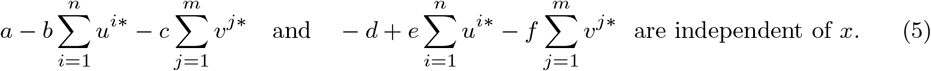

Unless otherwise specified, we always take 𝒮 as a nonempty subset of {1, …, *n*} and 𝒦 as a nonempty subset of {1, …, *m*}, and 𝒮^*c*^ = {1, …, *n*} \ 𝒮 and 𝒦^*c*^ = {1, …, *m*} \ 𝒦 in the remaining of this paper.

### 1.3 Main results

In competitive systems, species failing to form IFD may be invaded by rare exotic species with different dispersal strategies (Cantrell et al. 2021; Lam and Zhang 2025). We address such a problem for prey-predator system (1). The result implies the necessity of IFD even in the autonomous systems case of Cantrell and Cosner (2025). Without loss of generality, we illustrate it in the setting of *m* = *n* = 2. The result can be stated as follows.

#### Theorem 2

*Let m* = *n* = 2 *for system* (1). *Suppose that time-periodic distribution* (*u*^1*^, *v*^1*^) *is not an IFD. Then there exists a dispersal strategy* 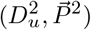 *or* 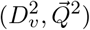 *such that* (*u*^1*^, 0, *v*^1*^, 0) *is unstable for system* (1).

This indicates that if a pair of resident prey and predator species do not form an IFD, then they can be invaded, i.e., any non-ideal free species distribution is not evolutionarily stable. One can similarly prove the instability of any non-IFD periodic solutions consisting of multiple prey-predator species.

Next, we generalize Theorem 1 to the time-periodic system (1) with arbitrarily many prey and predator species. It turns out that we need to solve an auxiliary non-autonomous predator-prey ODE system (see (7)). We will prove that if a 𝒮 - 𝒦 joint IFD exists, then under certain assumptions on the environmental coefficients 𝒜 := (*a, b, c, d, e, f*), it is stable in the evolutionary sense. To state the assumptions on 𝒜, we need to use time-periodic functions (*M*(*t*), *N*(*t*)), which can be defined entirely in terms of 𝒜. First we introduce the generalization of conditions **(Ha)-(Hb)**.

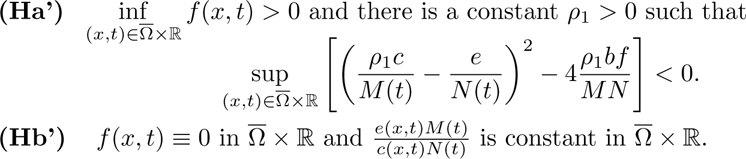

where (*M*(*t*), *N*(*t*)) are quantities derived entirely in terms of the coefficients 𝒜 := (*a, b, c, d, e, f*). The definition of of (*M*(*t*), *N*(*t*)) is contained in Subsection 2.2.

*Remark 3* If 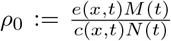 is constant in 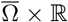 and inf *f >* 0, then **(Ha’)** holds for *ρ*_1_ sufficiently close to *ρ*_0_.

*Remark 4* When *a, b, c, d, e >* 0 and *f* ≥ 0 are time-independent, *M*(*t*) and *N*(*t*) are positive constants (see Appendix A). Then assumptions **(Ha’)-(Hb’)** reduce to **(Ha)-(Hb)**.

The following results show the evolutionary advantage of IFDs when the system is with or without self-limitation of the predator species (i.e, *f*(*x, t*) *>* 0 or *f*(*x, t*) ≡ 0 for all 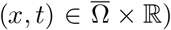. Specifically, the combination of species will be selected when this combination forms a unique IFD. This is the special case of Theorems 10 and 11 that are discussed in Sections 4 and 5, respectively.

#### Theorem 3

(With self-limitation) *Consider* 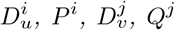 *and a, b, c, d, e, f satisfying* **(A)** *and* **(Ha’)**. *Suppose there exists a unique combination* 𝒮 ⊆ {1, …, *n*} *and* 𝒦 ⊆ {1, …, *m*} *such that the system* (1) *has a* 𝒮 ^*′*^*-*𝒦^*′*^ *joint IFD if and only if* (𝒮 ^*′*^, 𝒦^*′*^) = (𝒮, 𝒦). *Then for each positive solution* 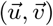 *to the initial value problem* (1), *we have*

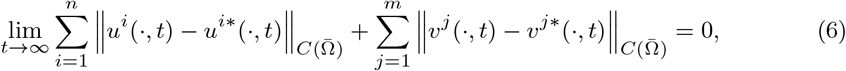

*where* 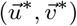 *is the* 𝒮 *-* 𝒦 *joint IFD*.

An immediate consequence of Theorem 3 is the following.

**Corollary 4** Under the assumptions of Theorem 3, suppose in addition that 𝒮 = {1} and 𝒦 = {1} are the unique subsets of prey-predator species that jointly support an IFD. Then

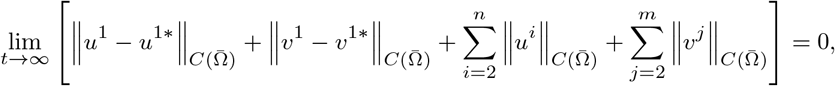

where 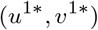 is the IFD.

#### Theorem 5

(Without self-limitation) *Let* 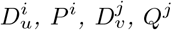 *and a, b, c, d, e, f satisfy* **(A)** *and* **(Hb’)**. *Suppose there exists a unique combination* 𝒮 ⊆ {1, …, *n*} *and* 𝒦 ⊆ {1, …, *m*} *such that the system* (1) *has a* 𝒮^*′*^*-* 𝒦^*′*^ *joint IFD if and only only if* (𝒮^*′*^, 𝒦^*′*^) = (𝒮, 𝒦). *Assume, in addition, that* (1) *has no nontrivial periodic solutions* 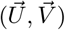 *such that U*^*i*^ ≡ 0 *for all i* ∈ 𝒮, *and*

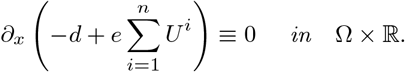

*Then for each positive solution of the system* (1), *the convergence* (6) *holds*.

### 1.4 Organization of the paper

In Section 2, we present some preliminary results such as the characterization of IFD in terms of (*M*(*t*), *N*(*t*), *K*^(1)^(*x, t*), *K*^(2)^(*x, t*)), which are quantities derived from the environmental coefficients. Section 3 is devoted to the existence of IFD, which is different from the case of temporally constant environments (Cantrell and Cosner 2025). The time-periodic environments may sometime contain “generalized sinks” that prevent the species from adopting an IFD. We give necessary and sufficient conditions for a given environment (Ω, *a, b, c, d, e, f*) to achieve an IFD. In Sections 4 and 5, we show the evolutionary stability of IFD with and without predator self-limitation, respectively. We prove Theorem 2 in Section 6. In Section 7, we demonstrate numerically that being able to approximate the conditions for IFD is utilitarian and confers advantage to the community of species, i.e. the evolutionary stability of IFD is in this sense robust. This complements and extends our analytical results. Finally, we discuss and interpret our results in the context of evolutionary community ecology in Section 8.

## 2 Preliminary

### 2.1 Well-posedness

Similar to Proposition 1 in Cantrell and Cosner (2018), we can transform system (1) into a reaction-diffusion-advection system with Robin boundary conditions, which admits a comparison principle (Lam and Lou 2022). Then, following standard theory (Lunardi 1995), we have the global existence and uniform boundedness of classical solutions to (1).

#### Theorem 6

*Suppose that* **(A)** *holds. Then system* (1) *with nonnegative, continuous initial data* 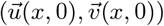 *has a global positive solution* 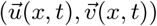, *and for any γ* ∈ (0, 1),

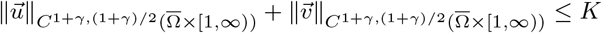

*for some constant K >* 0.

### 2.2 Definition of *M* (*t*), *N* (*t*) in terms of environmental coefficients

We will define the quantities (*M*(*t*), *N*(*t*)) as a positive time-periodic solution to the auxiliary ODE system

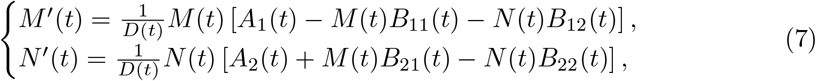

where *A*_*i*_, *B*_*ij*_ (with *B*_*ij*_ ≥ 0) are given entirely in terms of the environmental coefficients *a, b, c, d, e, f*:

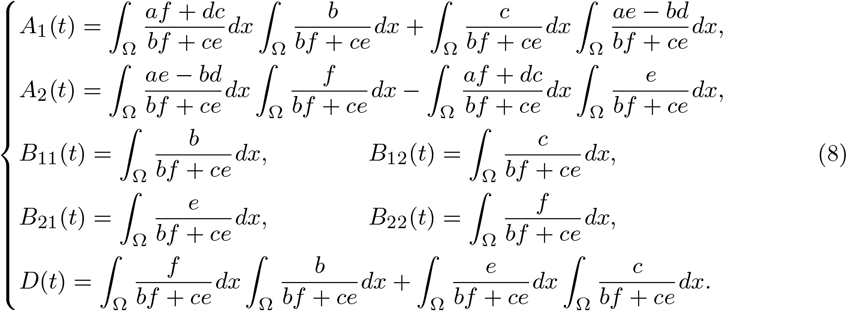

To motivate system (7), we begin by observing that whenever (*u*^*^, *v*^*^) is an IFD according to Definition 1, then ∫ _Ω_ *u*^*^(*x, t*) *dx*, ∫_Ω_ *v*^*^(*x, t*) *dx* is a positive solution of (7).

Indeed, denote

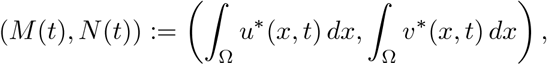

and observe that *a* − *bu*^*^ − *cv*^*^ and −*d* + *eu*^*^ − *fv*^*^ are independent of *x* and

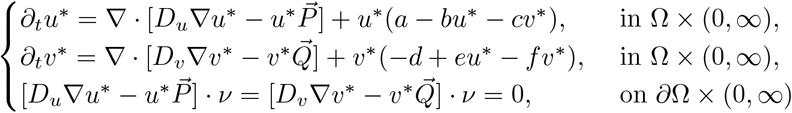

Integrating the above equation over Ω yields

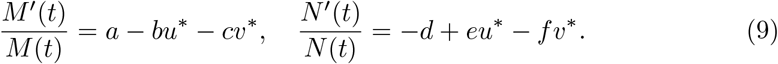

Multiply the first equation of (9) by *f* and the second one by *c*, subtract the resulting equations, and integrate over Ω to yield

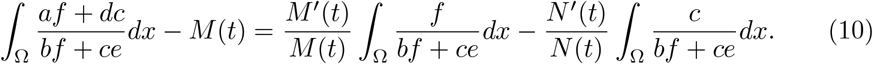

Repeat, but multiply the first equation of (9) by *e* and the second one by *b*, then we similarly get

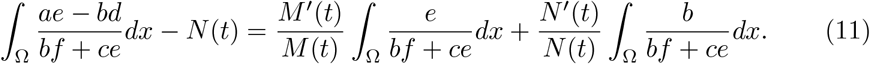

The equations (10) and (11), are equivalent to (7).

The existence of positive time-periodic (*M*(*t*), *N*(*t*)) is equivalent to the uniform persistence of both species (Smith and Thieme 2011), which has been studied extensively, e.g., see Teng (1999); Butler and Freedman (1981); Ding et al. (1995). To be more specific, we next give some sufficient conditions.

#### Lemma 7

System (7) has at least one positive *T*-periodic solution (*M*(*t*), *N*(*t*)) provided one of the following conditions holds:

i. *a, b, c, d, e, f >* 0 are time-independent and satisfy^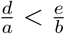^ for all 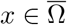.
ii. 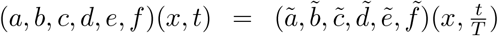, where 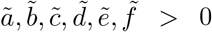 are given 1-periodic functions such that 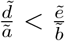 for all 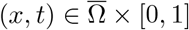 and *T* ≫ 1.
iii. *b, c, e, f >* 0 are independent of *x* and

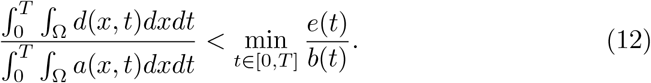
iv. System (1) has a 𝒮 − 𝒦 joint IFD.

Moreover, if one of assumptions **(Ha’)** and **(Hb’)** holds, then (*M*(*t*), *N*(*t*)) is unique.

*Proof* Indeed, if the system has a 𝒮 - 𝒦 joint IFD 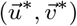, we can argue as in the beginning of Subsection (2.2) that 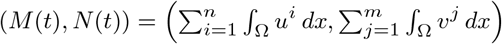 solves (7). The proofs of sufficient conditions (i)-(iii) are postponed to Appendix and that of (iv) is given in Theorem 9, and the uniqueness under **(Ha’)** (resp. **(Hb’)**) is a result of Theorem 10 (resp. Theorem 11). □

#### Definition of *K*^(1)^, *K*^(2)^ in terms of (*M* (*t*), *N* (*t*))

For a given positive solution (*M*(*t*), *N*(*t*)) of (7), we define 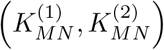 as follows

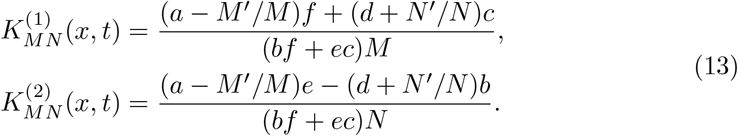

Note that we may sometimes suppress the subscript of 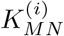 to simplify notations.

Since (*M*(*t*), *N*(*t*)) satisfies (10) and (11), it is easy to verify that *K*^(*i*)^(*x, t*) satisfies

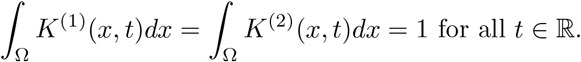

### 2.3 An auxiliary lemma

The following lemma plays an essential role in the evolutionary stability analysis. The proof of this lemma is the same as Lemma 1.2 in Lam and Zhang (2025), so we omit it.

#### Lemma 8

Suppose *d*(*x, t*) and 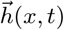 are smooth and *T*-periodic in *t*. Then

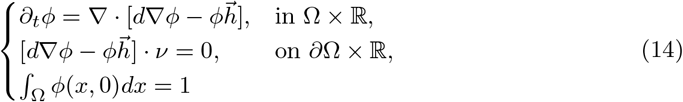

has a unique positive and *T*-periodic solution 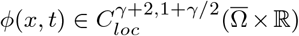 for any *γ* ∈ (0, 1).

Moreover, ∫_Ω_ *ϕ*(*x, t*)*dx* = 1 for all *t*.

#### Definition 3

For 1 ≤ *i* ≤ *n* (resp. 1 ≤ *j* ≤ *m*), we define 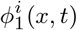 (resp. 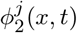) to be the unique positive *T*-periodic solution of (14) with 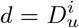 and 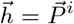 (resp. 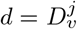 and 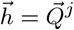).

## 3 Dispersal strategies that produce IFD

In this section, we investigate whether IFD is possible in a given environment. More precisely, for given environmental parameters (Ω, *a, b, c, d, e, f*), we determine whether there exist dispersal strategies 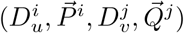, such that a 𝒮𝒦 joint IFD for system (1) can be achieved.

The following result gives the necessary and sufficient conditions for (1) to have an IFD including an arbitrary number of species.

### Theorem 9

*The following statements are equivalent:*

i. *System* (1) *has a* 𝒮 *-* 𝒦 *joint IFD* 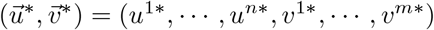.
ii. *System* (7) *has a positive solution* (*M*(*t*), *N*(*t*)) *and the corresponding K*^(1)^(*x, t*) *and K*^(2)^(*x, t*) *defined by* (13) *are positive*.

*Moreover*,

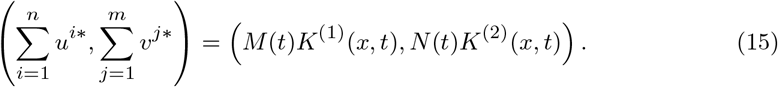

*Proof* We first prove “(i)⇒(ii)”. By Definition 2, 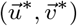 satisfies

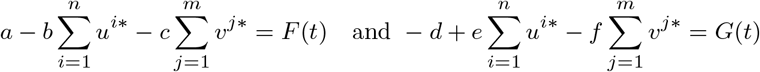

for *T*-periodic functions *F*(*t*) and *G*(*t*), and for all *i, j*,

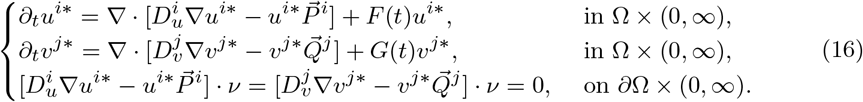

Define 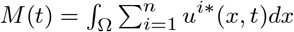 and 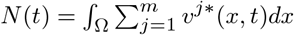. Then *M*(*t*), *N*(*t*) *>* 0 are *T*-periodic. Summing (16) over *i, j* and integrating over Ω yields

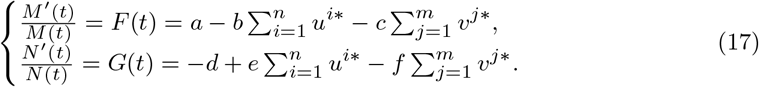

Using the similar arguments for (10) and (11), we deduce that (*M*(*t*), *N*(*t*)) is a positive solution of (7).

Note that *K*^(1)^(*x, t*), *K*^(2)^(*x, t*) defined by (13) satisfy

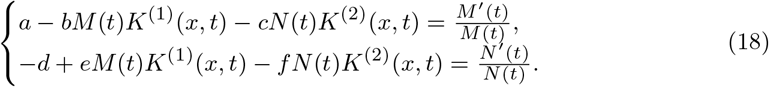

Subtracting (18) from (17) yields

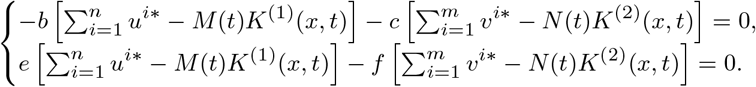

This proves (15), which implies the pointwise positivity of *K*^(1)^(*x, t*), *K*^(2)^(*x, t*).

We now prove “(ii)⇒(i)”. Suppose that (*M*(*t*), *N*(*t*)) is a positive and *T*-periodic solution of (7), and *K*^(1)^(*x, t*), *K*^(2)^(*x, t*) defined in terms of *M*(*t*), *N*(*t*) by (13) are positive. Then *M*(*t*)*K*^(1)^(*x, t*) and *N*(*t*)*K*^(2)^(*x, t*) are positive and *T*-periodic in *t*. By (18) and Definition 1, the distribution (*M*(*t*)*K*^(1)^(*x, t*), *M*(*t*)*K*^(2)^(*x, t*)) is an IFD.

It suffices to show (*M*(*t*)*K*^(1)^(*x, t*), *M*(*t*)*K*^(2)^(*x, t*)) is a solution to (1) with *n* = *m* = 1 by choosing suitable dispersal strategies 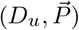 and 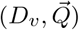. For given *D*_*u*_(*x, t*) and *D*_*v*_(*x, t*), take

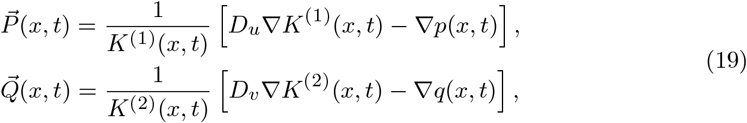

where *p*(*·, t*), *q*(*·, t*), for each *t* ∈ ℝ, uniquely solve

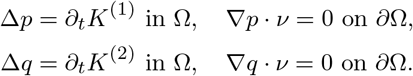

This and (19) imply that

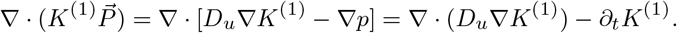

Then for *u* = *MK*^(1)^, a direct calculation yields

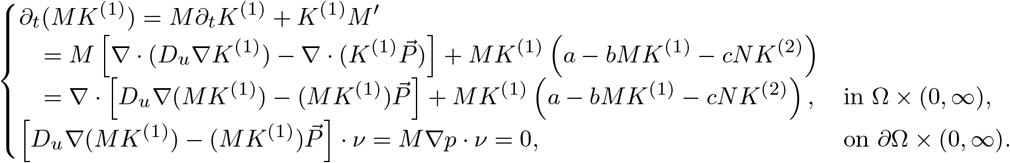

Similarly, *v* = *NK*^(2)^ satisfies

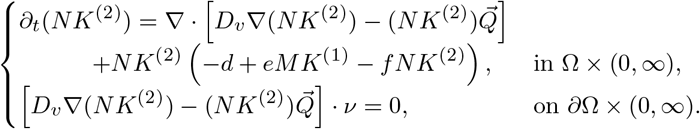

Therefore, (*M*(*t*)*K*^(1)^(*x, t*), *N*(*t*)*K*^(2)^(*x, t*)) is a solution of (1) with *m* = *n* = 1. □

Note that for general *n* and *m*, we can choose the dispersal strategies 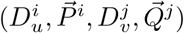 in the form of (19), such that system (1) has a 𝒮 - 𝒦 joint IFD 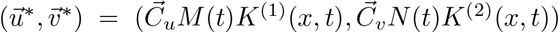, where 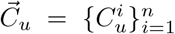 and 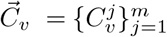 are constants satisfying 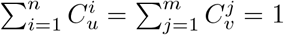.

## 4 Evolutionary stability with predator self-limitation

In this section, we establish that, in the presence of predator self-limitation, the total population of the prey-predator species approaches an IFD as *t* → ∞. In particular, any species that cannot participate in IFD is automatically excluded. Note that assertion (b) of the following implies Theorem 3.

### Theorem 10

(With self-limitation) *Suppose* 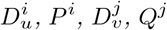 *and a, b, c, d, e, f satisfy* **(A)** *and* **(Ha’)**. *Then suppose system* (1) *has a* 𝒮 *-*𝒦 *joint IFD* 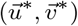. *Then for any solution* 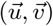 *to the initial value problem* (1) *with nontrivial nonnegative initial data, the following statements hold*.

a. *(The total distribution of the prey-predator species community approaches IFD) As the integer k* → +∞,

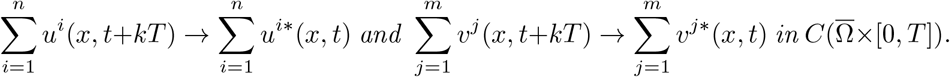
b. *(Only the unique combination of IFD species survives) In addition, suppose that for any nonempty sets* 𝒮^*′*^ ⊆ {1, …, *n*} *and* 𝒦^*′*^ ⊆ {1, …, *m*}, *system* (1) *has a* 𝒮^*′*^*-* 𝒦^*′*^ *joint IFD only when* 𝒮^*′*^ = 𝒮 *and* 𝒦^*′*^ = 𝒦. *Then, as t* → +∞,

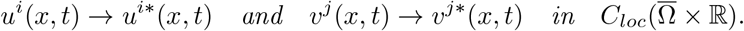

*Remark 5* Conclusion (b) of the Theorem 10 can be rephrased as follows: there exist nonnegative constants *p*^*i*^, *q*^*j*^ such that

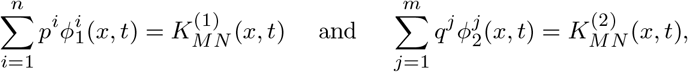

if and only if {*i* : *p*^*i*^ *>* 0} = 𝒮 and {*j* : *q*^*j*^ *>* 0} = 𝒦, where 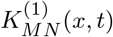 and 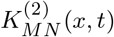 are given in (13), and 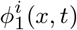 and 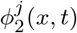 are given by Definition 3.

### 4.1 Proof of Theorems 10

*Proof* Throughout this proof, denote, for 1 ≤ *i* ≤ *n* and 1 ≤ *j* ≤ *m*,

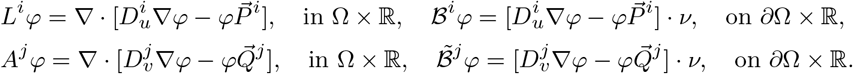

By Theorem 9, we obtain the existence of positive and *T*-periodic functions *M*(*t*), *N*(*t*), and that 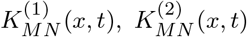 defined by (13) are positive. For convenience, we write *K*^(1)^(*x, t*), *K*^(2)^(*x, t*) in that which follows.

With the help of (17), *M*(*t*), *N*(*t*) and the 𝒮 -,𝒦 joint IFD, 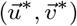 satisfy

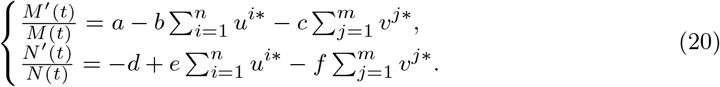

Hence, for all *i, j*,

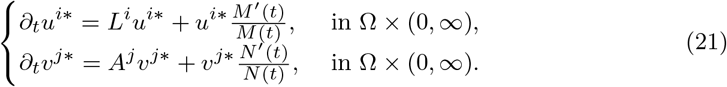

**Step 1**. Fix a solution 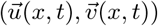 of system (1) with {*u*^*i*^}_*i*∈𝒮_ and {*v*^*j*^}_*j*∈𝒦_ being componentwise positive, define

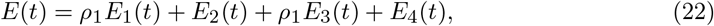

where *ρ*_1_ *>* 0 is given in **(Ha’)** and

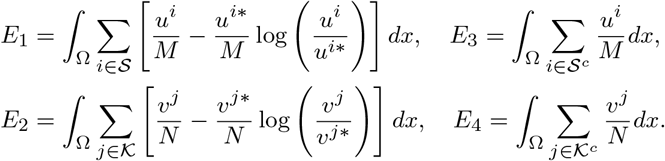

Thanks to the no-flux boundary conditions, we calculate that

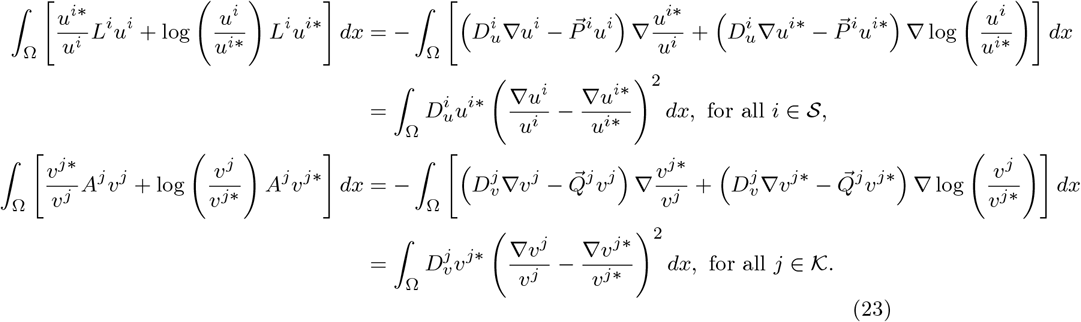

Moreover, we use (20) to deduce the following identities:

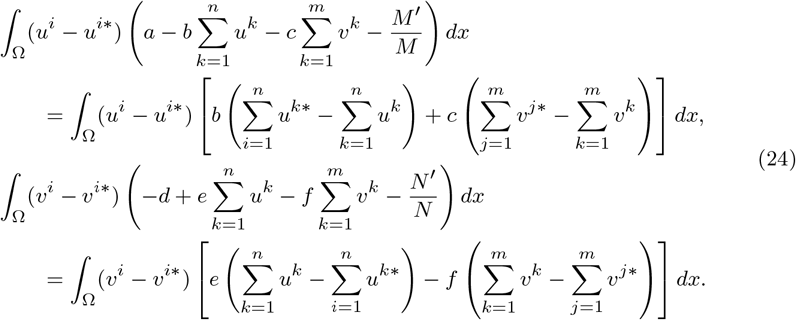

Now some calculations yield that

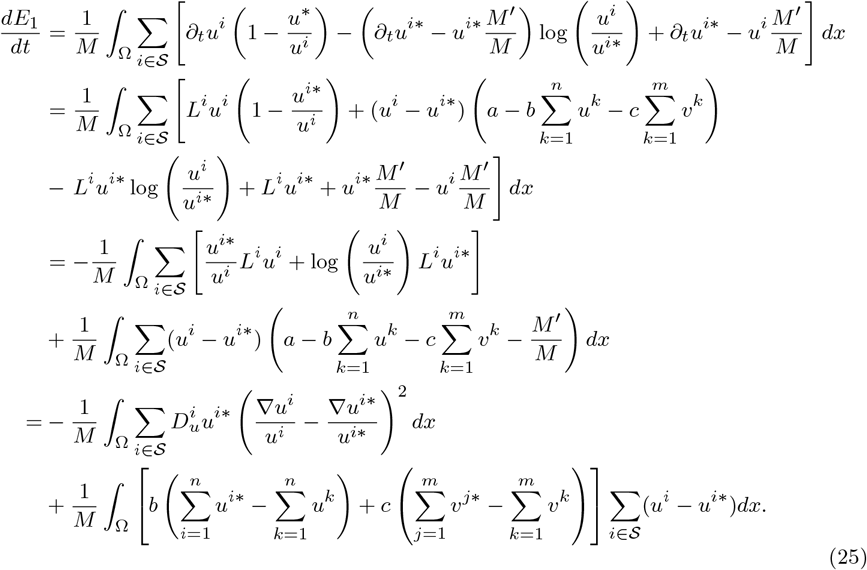

An analogous calculation gives

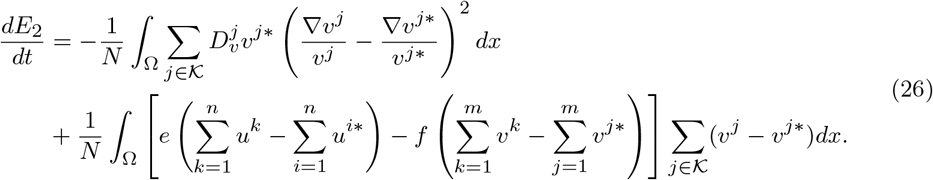

Again apply the no-flux boundary conditions and (20) to obtain that

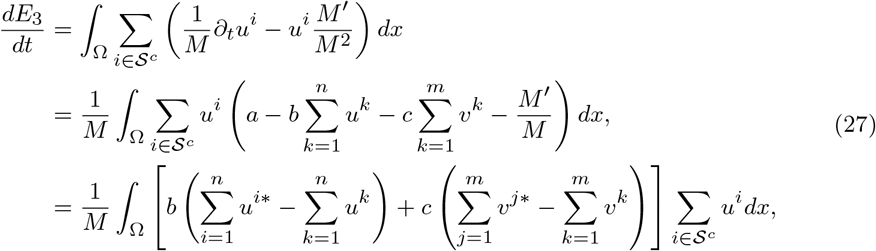

and

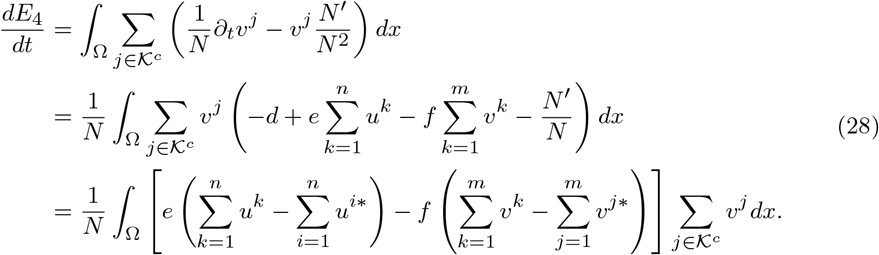

Substituting (25), (26), (27) and (28) into (22), we derive that

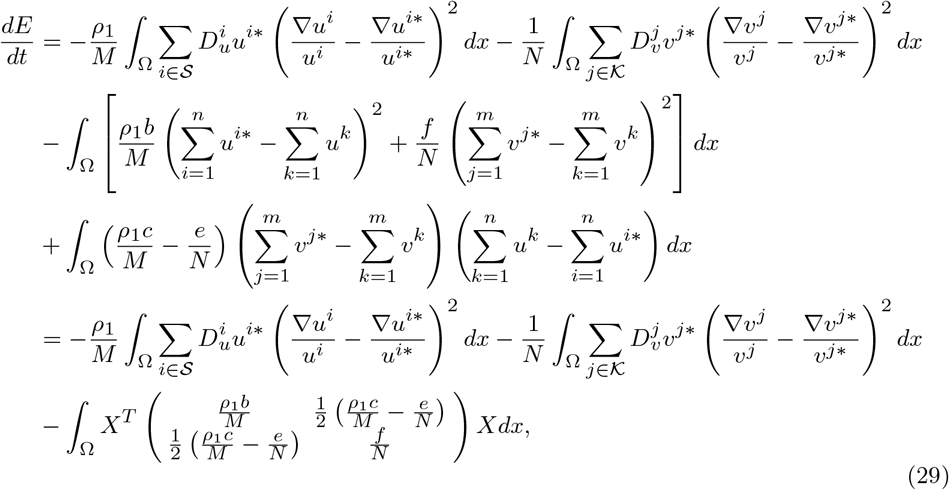

where 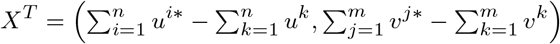.

Thanks to **(Ha’)**, the matrix

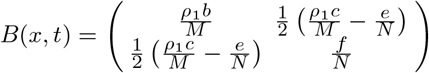

is positive definite for any 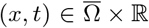, and hence there exists constant *K >* 0 such that for any 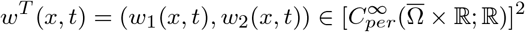,

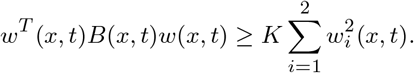

Therefore,

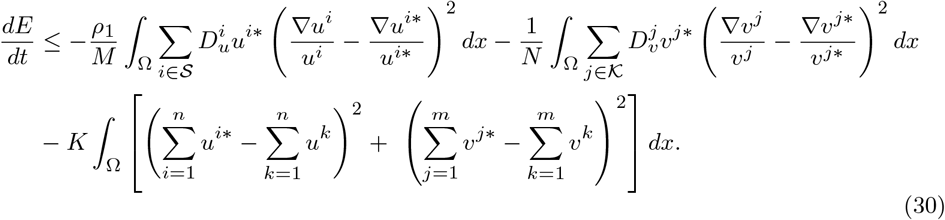

**Step 2**. Denote

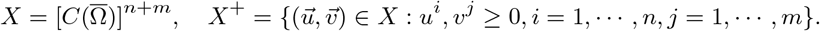

We note that the solution semiflow of system (1) is compact for any *t >* 0 and it is point dissipative by Theorem 6. It then follows from Theorem 1.1.2 (Zhao 2017) that system (1) has a global compact attractor 𝒜 ⊆ *X*^+^. For any solution 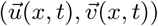 to (1) in Γ, LaSalle’s invariance principle (Zhao 2017, Theorem 1.1.1) implies that its *ω*-limit set is contained in ℳ^*′*^(*t*), the largest invariant subset of

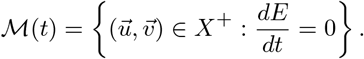

Here if we let 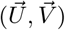 be a positive solution of (1) and suppose there are integers *N*_*k*_ → ∞ such that

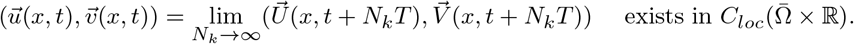

Then 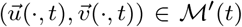 for all *t* ∈ ℝ. In the following, we abuse notations slightly to define 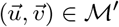 to mean 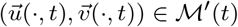 for all *t* ∈ ℝ.

We need to deduce properties of ℳ^*′*^ to prove assertions (a) and (b). In view of (30), 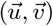 is an entire solution belonging to ℳ if and only if

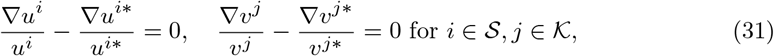

and

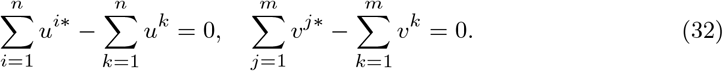

Clearly, (32) implies statement (a).

**Claim**. The function pair (*M*(*t*), *N*(*t*)) such that 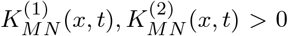 is unique.

This claim implies the uniqueness in Theorem 9. Otherwise, suppose that there exists another 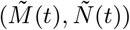 with 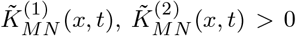. Without loss of generality, we consider the case *n* = 2, *m* = 2 and write

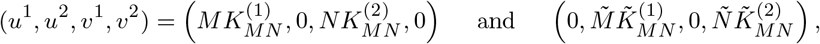

where both are IFDs. Then using (32) and integrating over Ω implies 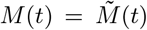 and 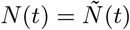.

Next, we assume there is a unique combination 𝒮 - 𝒦such that (1) achieves a 𝒮 - 𝒦joint IFD, and proceed to prove (b). By (32) and (20), the solution 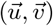 in the invariant set ℳ satisfies

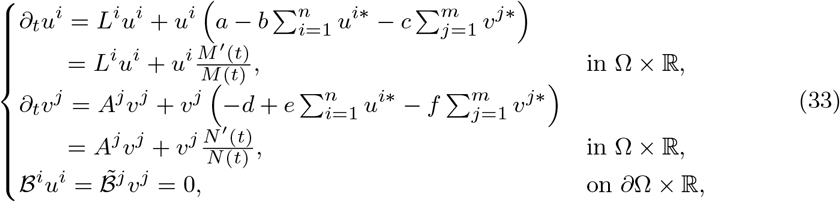

for all 1 ≤ *i* ≤ *n*, 1 ≤ *j* ≤ *m*.

**Step 3**. We claim that 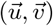 is *T*-periodic. By Lemma 8, there exist nonnegative constants 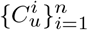 and 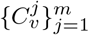 such that 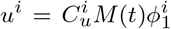 and 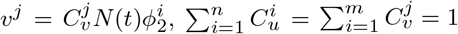, where 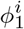 and 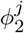 are given in Definition 3. Thus, 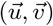 is *T*-periodic.

**Step 4**. We claim that, for every *i* ∈ 𝒮 (resp. *j* ∈ 𝒦), there exists a nonnegative constant 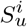 (resp. 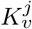) such that 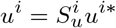 (resp. 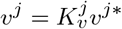). Upon combining with (32), we have

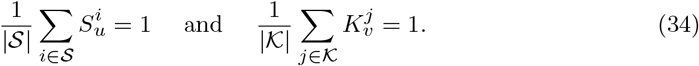

Indeed, (31) implies 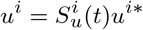 for each *i* ∈ 𝒮. Moreover, 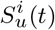 is *C*^1^ (since *u*^*i*^ and *u*^*i*\*^ are both *C*^1^ in *t*). Substituting 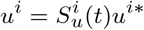 into *u*^*i*^-equation of (33) leads to 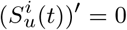. Thus, 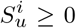 is a constant. Similarly, we have 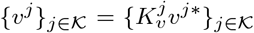 for some constants 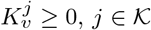. It suffices to prove the following claim, so that the invariant set ℳ^*′*^ consists of a single entire solution.

**Step 5**. We claim that 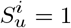 for all *i* ∈ 𝒮 and 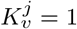 for all *j* ∈ 𝒦.

Suppose, on the contrary, that 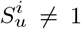 for some *i* ∈ 𝒮. (The case 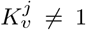 for some *j* ∈ 𝒦 can be treated in a similar manner.) Then by (34), there exists *i*_0_ ∈ 𝒮 such that 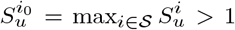. Now, if we set 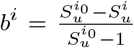 for *i* ∈ 𝒮 and *b*^*i*^ = 0 for *i* ∈ 𝒮 ^*c*^, then clearly 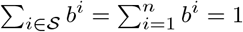. Define

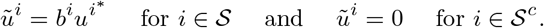

Then it is clear that 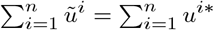 and that 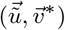 gives a 𝒮^*′*^ − 𝒦 joint IFD where 𝒮^*′*^ is a proper subset of 𝒮. This is a contradiction, so 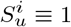 for all *i* ∈ 𝒮. A similar contradiction can be reached in 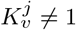 for some *j* ∈ 𝒦. □

## 5 Evolutionary stability without predator self-limitation

Now we consider the evolutionary stability of ideal dispersal strategies in the more delicate case where there is no self-limitation of the predator species. Theorem 5 is stated as assertion (b) in the following theorem.

### Theorem 11

(Without self-limitation) *Suppose* 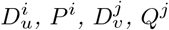 *and a, b, c, d, e, f satisfy* (**A**) *and* **(Hb’)**. *Then suppose that the system* (1) *has a* 𝒮 *-*𝒦 *joint IFD* 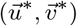. *Then for each solution* 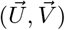 *initiating from any nontrivial and nonnegative data* 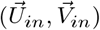, *the orbit*

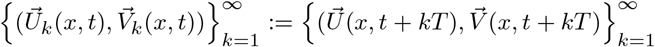

*has compact closure in* 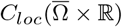, *which we denote by* 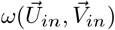.^1^

(a) *Any entire solution* 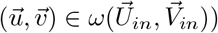 *(which is defined in* Ω *×* ℝ*) satisfies*

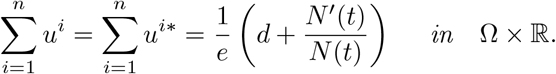

*Moreover, exactly one of the following holds:*

i. *The total distribution of* 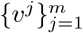 *is IFD, i.e*.

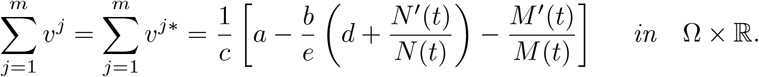
ii. *n >* 1 *and m >* 1 *and none of the species* {*u*^*i*^}_*i*∈𝒮_ *persists, i*.*e*.

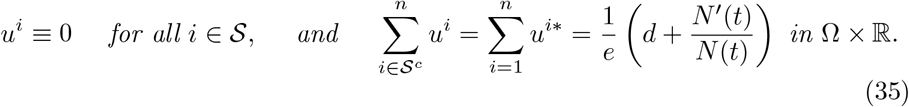

*Moreover, there exist nonnegative constants* 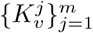 *such that*

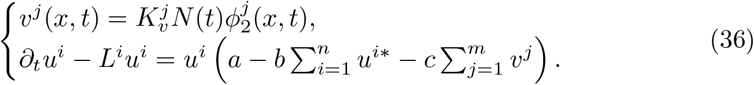

(b)*Suppose* 𝒮 − 𝒦 *is the unique combination that supports a joint IFD, and at least one of the following conditions holds:*

i. *n* = 1;
ii. *m* = 1;
iii. *n >* 1, *m >* 1, *and no subsets* 𝒮^*′*^ ⊆ 𝒮^*c*^ *and* 𝒦^*′*^ ⊆ {1, …, *m*} *can support a positive periodic solution* 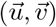 *such that* 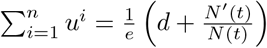.

*Then the following convergence holds uniformly in x* ∈ Ω:

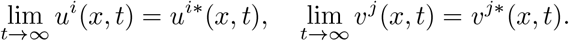

As an application, we derive the following exclusion principle when a single prey-predator pair forms an IFD.

**Corollary 12** Let 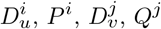 and *a, b, c, d, e, f* satisfy **(A)** and **(Hb’)**. Suppose the following two conditions hold simultaneously:

i. *n* = 1 or *m* = 1;
ii. 𝒮 = {1} and 𝒦= {1} are the unique subsets of prey-predator species that jointly support an IFD.

Then as *t* → ∞, the following convergences hold uniformly in *x* ∈ Ω:

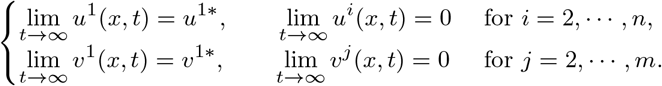

*Remark 6* Conditions (ii) of the above corollary can be rephrased as follows: suppose there exist nonnegative constants *p*^*i*^, *q*^*j*^ such that

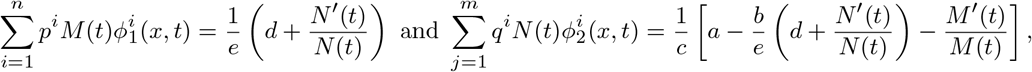

then {*i* : *p*^*i*^ *>* 0} = {1} and {*j* : *q*^*j*^ *>* 0} = {1}, where *ϕ*^*i*^ and *ϕ*^*j*^ are given in Definition 3.

### 5.1 Proof of Theorem 11

*Proof* By Theorem 9 and its proof, there exist positive *T*-periodic functions *M*(*t*), *N*(*t*) such that *K*^(1)^(*x, t*), *K*^(2)^(*x, t*) defined by (13) are positive, and the 𝒮 - 𝒦 joint IFD 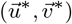 satisfies

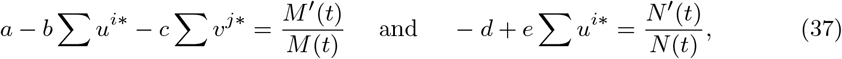

which implies that

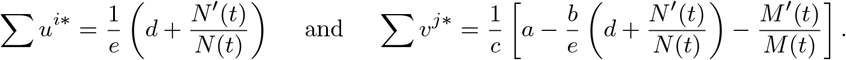

When **(Hb’)** is satisfied, set *f*(*x, t*) ≡ 0 and substitute *ρ*_2_ for *ρ*_1_ in (29) to derive

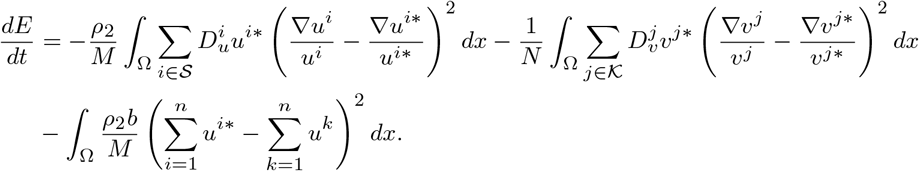

It follows that

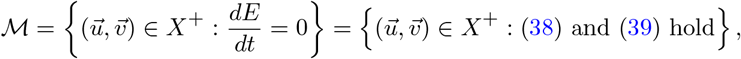

where

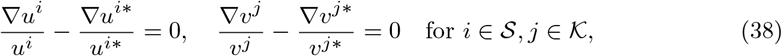

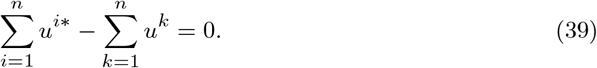

This establishes (11).

Suppose there exist alternative 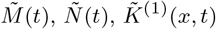 and 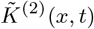. Let

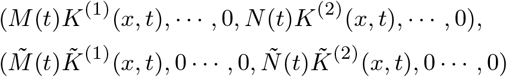

be two solutions of (1), then we will see that 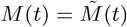 by integrating (39) over Ω, whereas the uniqueness of *N*(*t*) follows from the first equation of (7). This proves the uniqueness of (*M*(*t*), *N*(*t*))in Theorem 9.

We further get from (39) that the solution 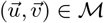 satisfies that

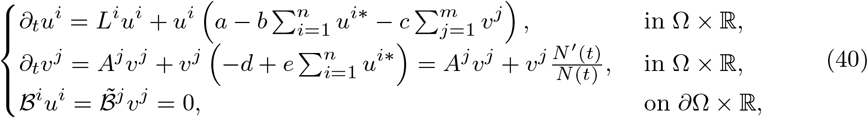

where 1 ≤ *i* ≤ *n*, 1 ≤ *j* ≤ *m*.

**Claim**. The solution 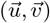 of (40) is *T*-periodic. Combining the *v*^j^-equation in (40) and Lemma 8, there exist nonnegative constants 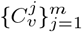 such that 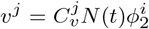, where 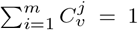 and 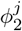 is given in Definition 3. Thus, 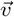 is *T*-periodic. Substituting 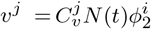 into *u*^*i*^-equation, and again using Lemma 8 to see *u*^*i*^ is *T*-periodic.

Next, we establish the dichotomy in assertion (a). We divide the proof into the following cases: (I) *u*^*i*^ ≠ 0 for some *i* ∈ 𝒮; (II) *n* = 1 (there is only one prey species); (III) *m* = 1 (there is only one predator species); (IV) *u*^*i*^ ≡ 0 for all *i* ∈ 𝒮. We will show that in cases (I), (II) and (III), the alternative (i) holds, and that (ii) holds in case (IV).

**Case (I)**. Suppose *u*^*i*^ ≢ 0 for some *i* ∈ 𝒮, then (38) implies that

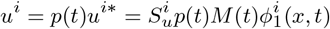 with some positive, periodic function *p*(*t*) and constant 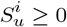.

Substituting into the *i*-th equation in (40), we see if 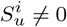 (i.e., *u*^*i*^ ≠ 0), then

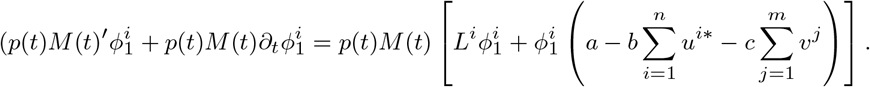

By Definition 3 and *p*(*t*) *>* 0, we deduce that 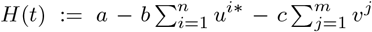 is independent of *x*.

We claim that 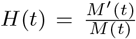. Indeed, integrating the equation of *u*^*i*^ in (40) in Ω and summing over 1 ≤ *i* ≤ *n*, we have

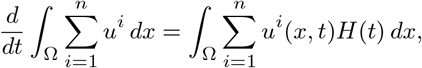

and we conclude the claim by (39) and 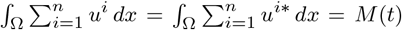. This, in turn, determines the sum 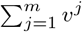 uniquely:

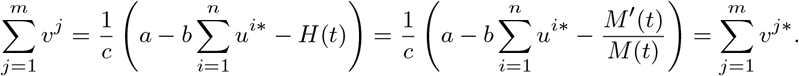

Hence, we showed that (i) holds if *u*^*i*^ ≢ 0 for some *i* ∈ 𝒮.

**Case (II)**. If *n* = 1, then we must have 𝒮 = {1} and *u*^1^ *?*≡ 0 thanks to (39). Hence *n* = 1 implies alterative (i) holds.

**Case (III)**. Assume *m* = 1. In this case, we must have 𝒦 = {1}, so that 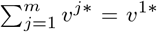 and 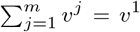. Integrating the equations of *u*^*i*\*^ in Ω and adding over 1 ≤ *i* ≤ *n*, we derive that

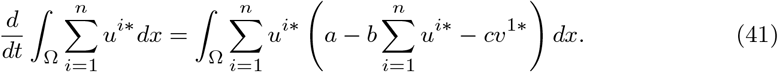

By the equation of *v*^1^ in (40) and that of *v*^1*^, we see that *v*^1^ = *qv*^1*^ for some constant *q >* 0.

Now, we integrate the equations of *u*^*i*^ in Ω and add over 1 ≤ *i* ≤ *n*, and obtain

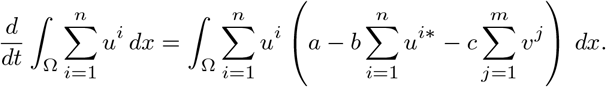

Using 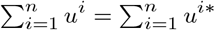 by (39) and 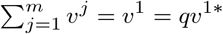, we have

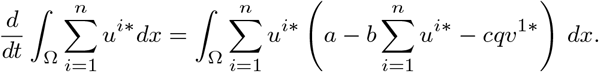

Subtract this from (41) to show 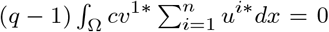, which implies that *q* = 1, and hence (i) holds.

**Case (IV)**. Finally, we assume that *u*^*i*^ ≡ 0 for all *i* ∈ 𝒮 (which implies *m >* 1 and *n >* 1). We need to deduce (ii). Substituting *u* 0 into (39), we deduce (35). Since *v*^j^ is a positive solution to a linear equation in (40), it follows by the Lemma 8 that 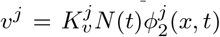 for some constant 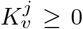 and is therefore *T*-periodic, where 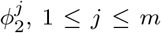, are given in Definition 3. Substituting back to (40) proves the conclusion of (ii). This finishes the proof of assertion (a).

Next, we prove assertion (b). Under the additional assumption, we conclude from part (a) that

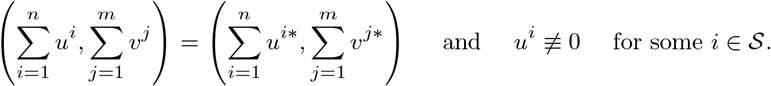

Now, 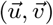 is a positive entire solution satisfying the system (33) with *f* ≡ 0. From this point, we can repeat Step 3, Step 4, and Step 5 in the proof of Theorem 10. This concludes the proof. □

## 6 Proof of Theorem 2

*The proof of Theorem 2* The eigenvalue problem corresponding to the linearized system of (1) with *m* = *n* = 2 at (*u*^1*^, 0, *v*^1*^, 0) is

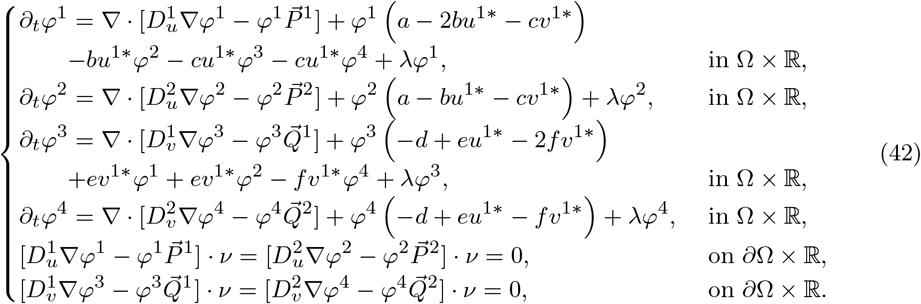

It suffices to prove that problem (42) has at least one eigenvalue with negative real part when (*u*^1*^, *v*^1*^) is not an IFD, where (*u*^1*^, *v*^1*^) satisfies

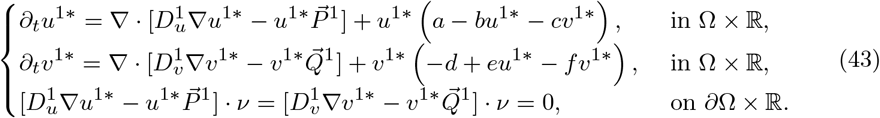

**Case (a)**. The following problem (44) has at least one eigenvalue with negative real part:

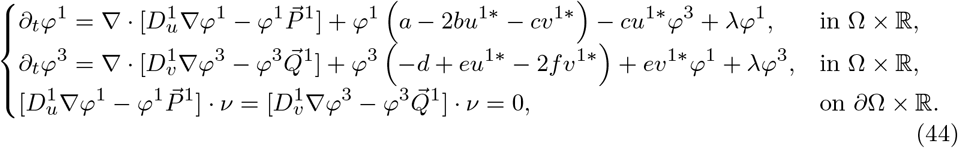

We prove the conclusion by choosing eigenfunction as (*φ*^1^, 0, *φ*^3^, 0), where (*φ*^1^, *φ*^3^) are the eigenfunction of (44) corresponding to the eigenvalue with negative real part.

**Case (b)**. All eigenvalues of problem (44) have nonnegative real parts. Since (*u*^1*^, *v*^1*^) is not an IFD of (43), we have

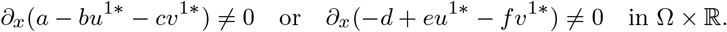

**Case (i)**. If *∂*_*x*_(*a* − *bu*^1*^ − *cv*^1*^) ≠ 0, integrate the first equation of (43) over Ω, then

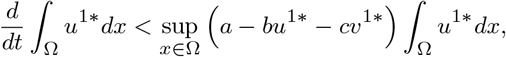

which implies that

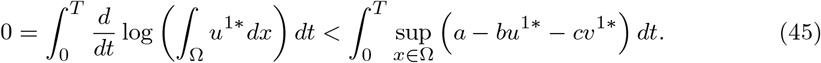

By (45) and Lemma B.1 (Cantrell et al. 2021) (i.e, 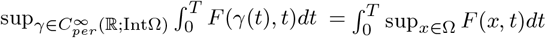 for all 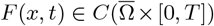), there exists 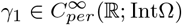 such that

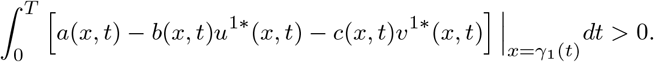

Thus, using the arguments in Theorem 4.2 (Cantrell et al. 2021), for given 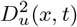 satisfying (**A**), we can choose

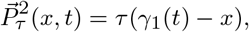

then lim sup_*τ* →∞_ *λ*_1_ *<* 0, where *λ*_1_ is the principal eigenvalue of the following problem

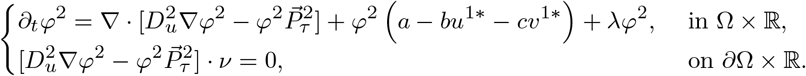

Denote the corresponding eigenfunction to *λ*_1_ as *φ*^2*^, and choose *φ*^4*^ = 0. Then substitute (*λ*_1_, *φ*^2*^, *φ*^4*^) into the first and third equations of (42) and obtain

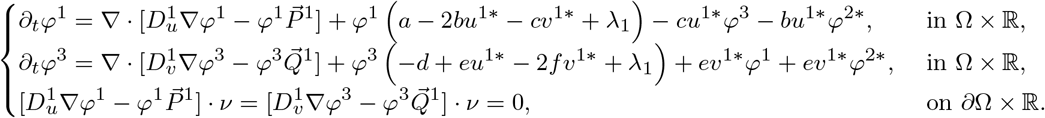

Observe that *λ*_1_ *<* 0 is not an eigenvalue of (44), so that the above system has a unique solution (*φ*^1*^, *φ*^3*^).

**Case (ii)**. If *∂*_*x*_(*a* − *bu*^1*^ − *cv*^1*^) ≡ 0 and *∂*_*x*_(−*d* + *eu*^1*^ − *fv*^1*^) ≠ 0, then we get

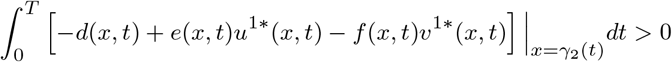

holds for some 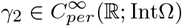. Moreover, fix 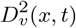 satisfying **(A)**, choose

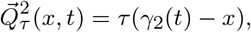

then lim sup_*τ* →∞_ *λ*_1_ *<* 0, where *λ*_1_ is the principal eigenvalue of the following problem

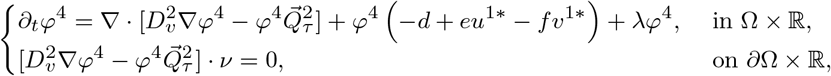

and the associated eigenfunction is *φ*^4*^. In a similar manner to **Case (i)**, we can prove that (42) has an eigenvalue *λ*_1_ with lim sup_*τ*_ *λ*_1_ *<* 0 and the associated eigenfunction is (*φ*^1*^, 0, *φ*^3*^, *φ*^4*^). □

## 7 Convergence stability of approximate IFD

We now conduct numerical simulations to show that in lieu perfectly achieving IFD, being able to approximate the conditions for IFD is utilitarian. This corresponds to the concept of convergence stability (Christiansen 1991; Eshel et al. 1997), means that the closer a (collection of) species is to IFD, then it has more competitive advantage against species that are further away from IFD. In particular, this suggest competitive advantage when IFD is approximately achieved. According to the basic timescale separation assumption of adaptive dynamics, upon repeated introduction of mutant prey/predator species, the total population evolves closer and closer towards achieving an IFD.

To study convergence stability, it is necessary to introduce a *metric* to quantify the deviation of a given distribution (in space-time coordinates) from IFD. We fix our discussion by defining a function *G* on the set Λ := {(𝒮^*′*^, 𝒦^*′*^) : ∅ = 𝒦^*′*^ ⊆ {1, …, *m*}} as

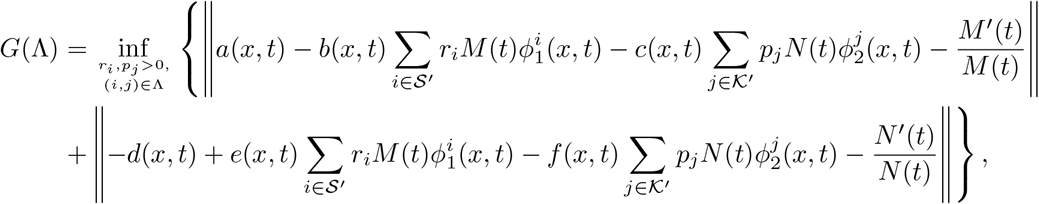

where 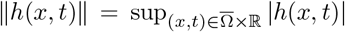, and 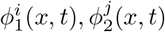 are defined by Definition 3. Observe that *G*(Λ) measures the deviation of the distribution of species from achieving IFD. Namely, *G*(Λ) ≥ 0 and *G*(Λ) = 0 if and only if Λ forms an IFD.

For each subset of species, we let the species to reach a terminal, stable time-periodic solution Λ, then we compute the corresponding value of *G*(Λ). To show convergence stability, we demonstrate with our numerical example that species subsets while distribution is closest to an IFD, i.e. that generats a smaller value of *G*(Λ), will have a competitive advantage.

### With self-limitation

Consider *n* = 2, *m* = 2, Ω = (0, 3*π*) and *T* = 24. In the case of *f*(*x, t*) *>* 0 for all (*x, t*) ∈ [0, 3*π*] *×* ℝ, take the following parameters:

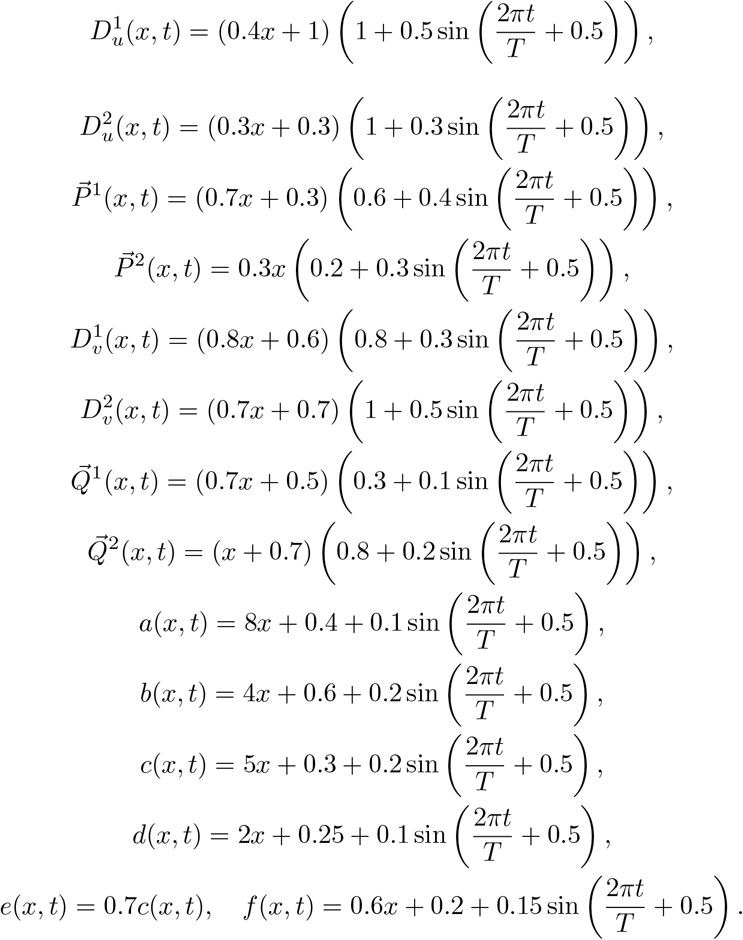

Under our choice o parameters, system (7) admits a positive solution (*M*(*t*), *N*(*t*)) and the corresponding *K*^(1)^(*x, t*) *>* 0, *K*^(2)^(*x, t*) *>* 0 (Fig. 1). By Theorem 9, system (1) with *n* = *m* = 2 has an *𝒮*-*𝒦* joint IFD. The values of *G*(Λ) are computed as follows:

**Fig. 1.**
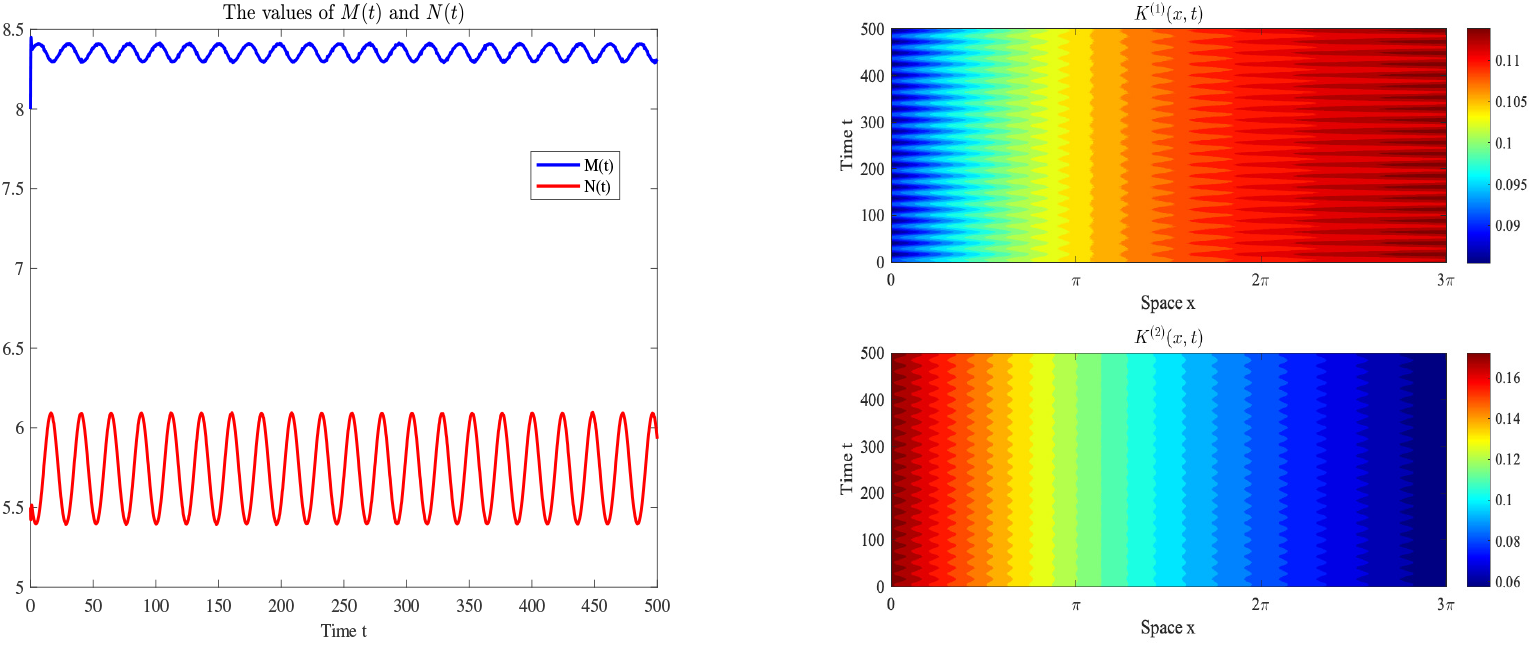
The graphs of *M*(*t*), *N*(*t*) and *K*^*(1)*^(*x, t*), *K*^*(2)*^(*x, t*).

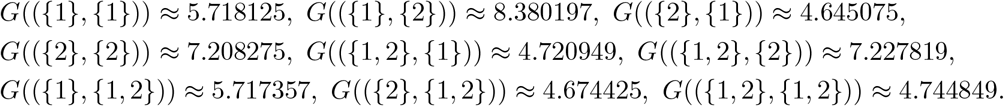

Obviously, the function *G* attains its unique minimum at ({2}, {1}). This implies that the species combination of the second prey and the first predator is closest to one IFD, and is more likely to be selected, which is confirmed by Fig. 2.

**Fig. 2.**
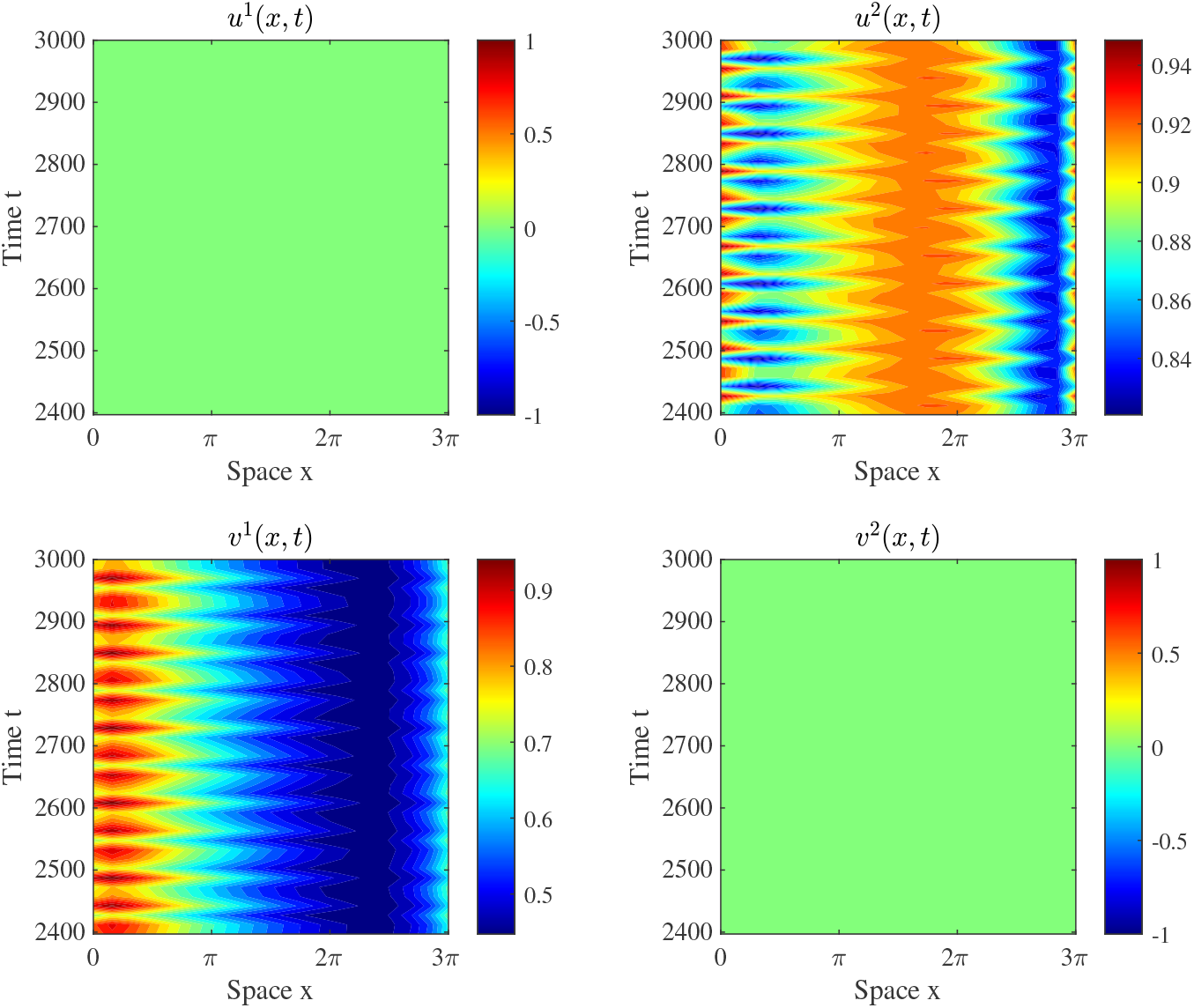
The solution of (1) with two prey species and two predator species, where parameters are given at the beginning of the section.

We perturb the dispersal strategy of the second prey species by modifying its directed movement as

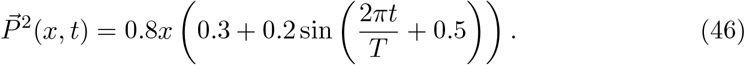

Despite this perturbation, the functional *G* maintains its unique minimum at ({2}, {1}), and numerical simulation also shows the coexistence of the second prey and first predator species (Fig. 3). This demonstrates that dispersal strategies that generate an approximation of IFD confer a competition advantage and it is not easily invaded.

**Fig. 3.**
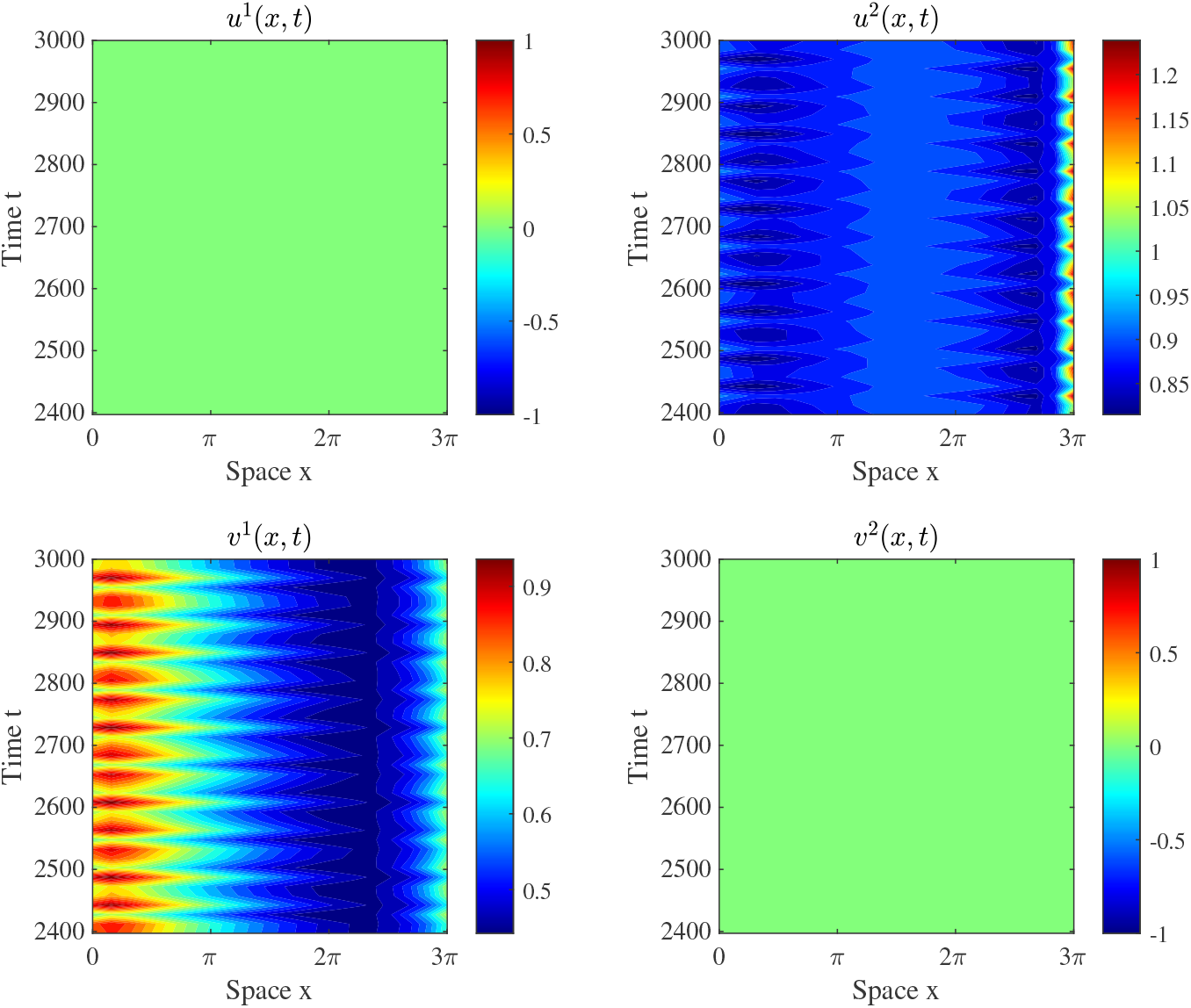
The solution of (1) with two prey species and two predator species, where parameters are given in the beginning of Section 7 except for 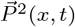.

### Without self-limitation

We now turn to the case of *f*(*x, t*) ≡ 0 for all (*x, t*) ∈ [0, 3*π*] *×* ℝ. Take the following parameters:

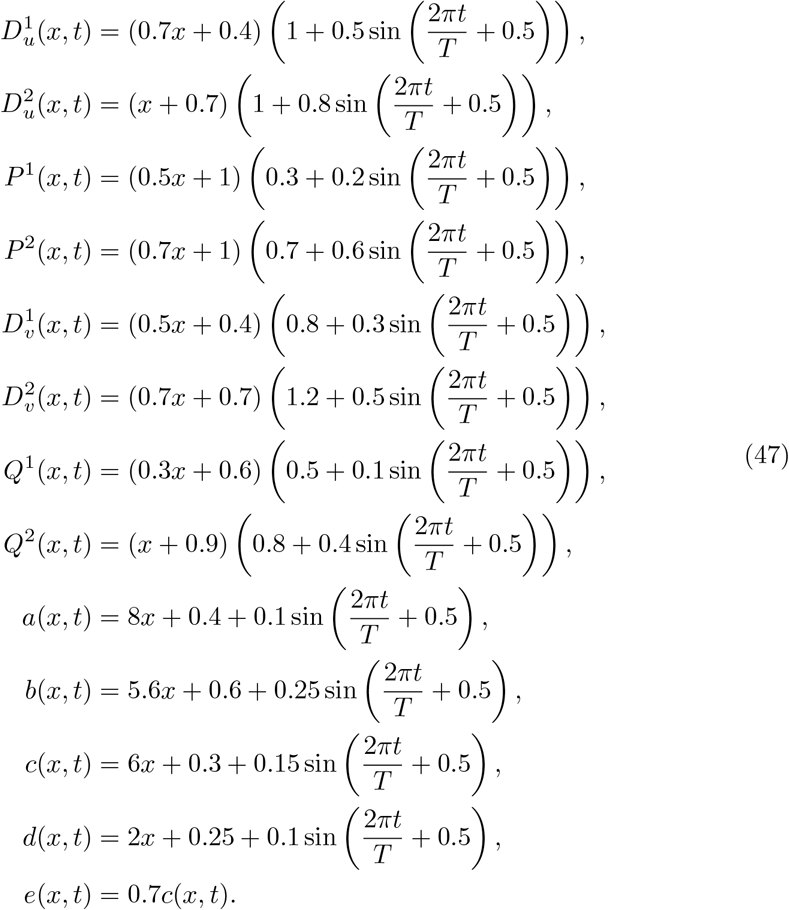

It can be verified that *M*(*t*) *>* 0, *N*(*t*) *>* 0, *K*^(1)^(*x, t*) *>* 0, *K*^(2)^(*x, t*) *>* 0 through numerical analysis, see Fig. 4. It follows from Theorem 9 that system (1) with *n* = *m* = 2 admits IFD. Further computations yield

**Fig. 4.**
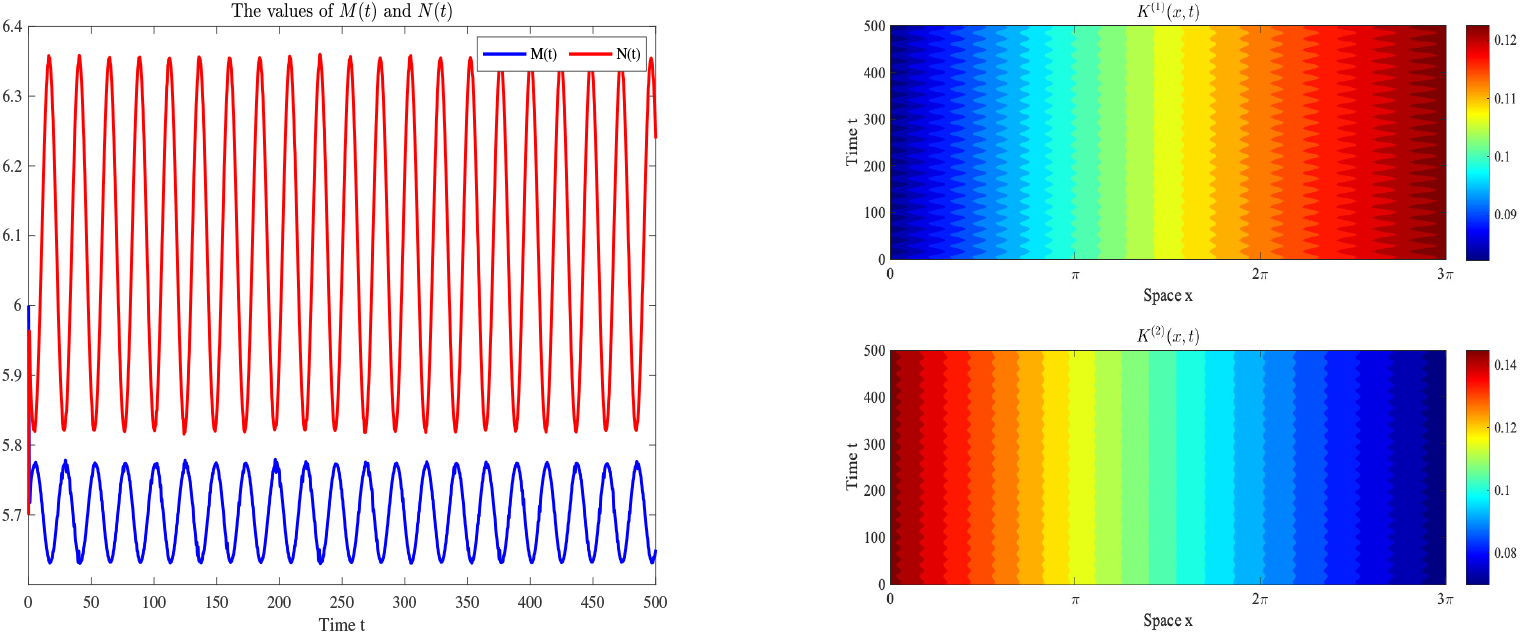
The graphs of *M*(*t*), *N*(*t*) and *K*^*(1)*^(*x, t*), *K*^*(2)*^(*x, t*).

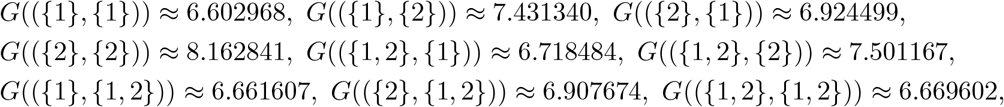

and hence 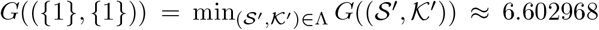. Meanwhile, Fig. 5 shows that the first prey and the first predator species prevail in the predation-competition interactions, which is consistent with that the combination with smallest *G* wins.

**Fig. 5.**
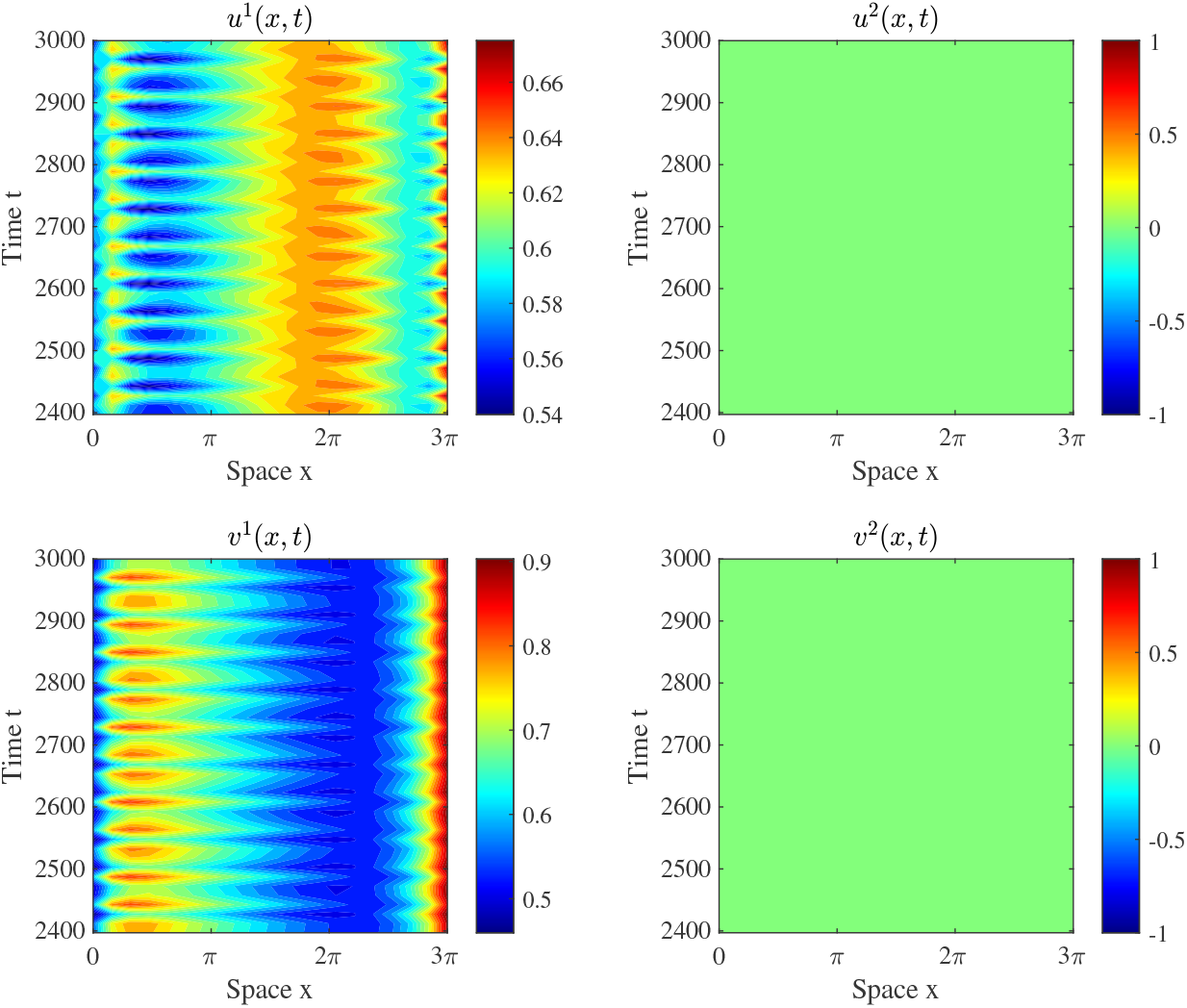
The solution of (1) with two prey species and two predator species, where parameters are given in (47).

### Invasion of non-ideal free dispersal strategies

In what follows, we show that the prey and predator species not jointly forming IFD will be invaded. Take the following parameters for system (43):

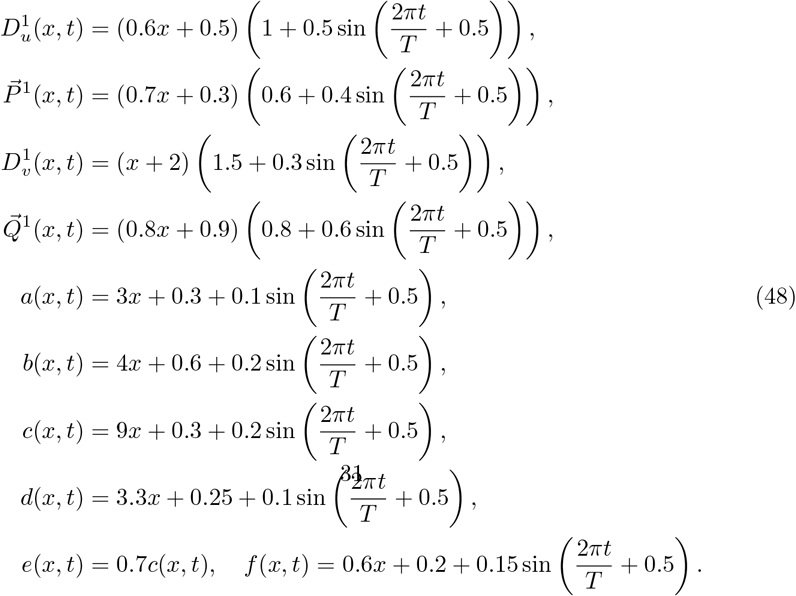

Then system (43) has a stable positive periodic solution (*u*^1*^, *v*^1*^) (Fig. 6(the left two panels)). Since *K*^(2)^(*x, t*) *<* 0 for some (*x, t*) ∈ [0, 3*π*] *×* [0, 500] (Fig. 6(the right two panels)), we see from Theorem 9 that (*u*^1*^, *v*^1*^) is not an IFD of (43). Introduce a second prey-predator pair (*u*^2^, *v*^2^) to system (43) with dispersal strategies:

**Fig. 6.**
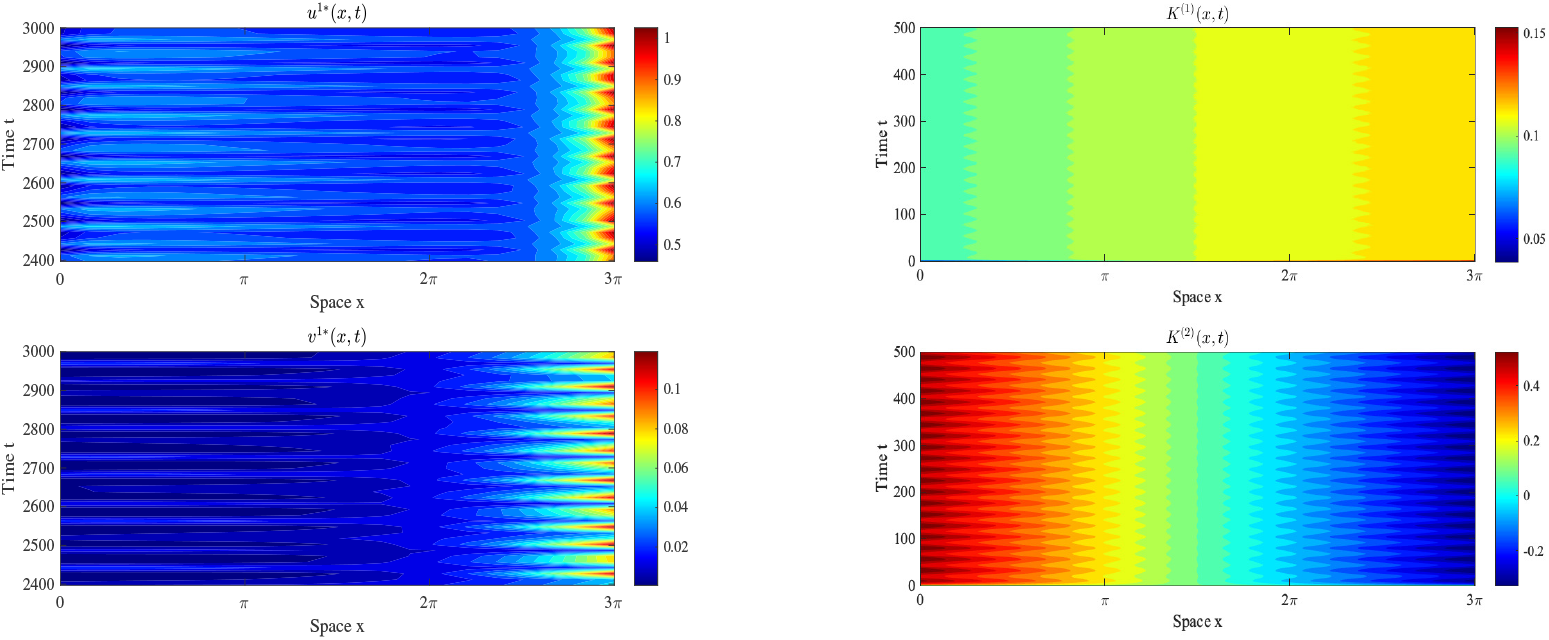
The graphs of (*u*^*1*\*^(*x, t*), *v*^*1*\*^(*x, t*)) and *K*^*(1)*^(*x, t*), *K*^*(2)*^(*x, t*).

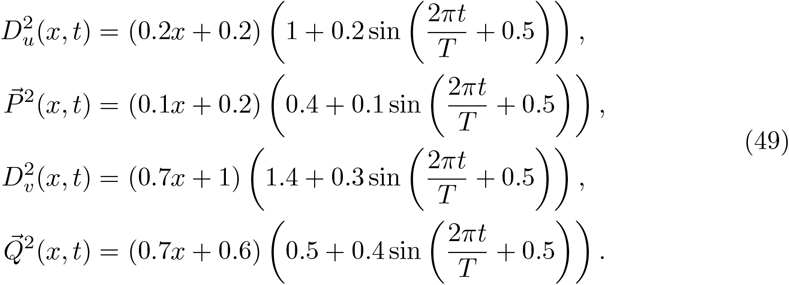

Numerical simulation demonstrates that species (*u*^2^, *v*^2^) drive (*u*^1^, *v*^1^) to extinction (see Fig. 7).

**Fig. 7.**
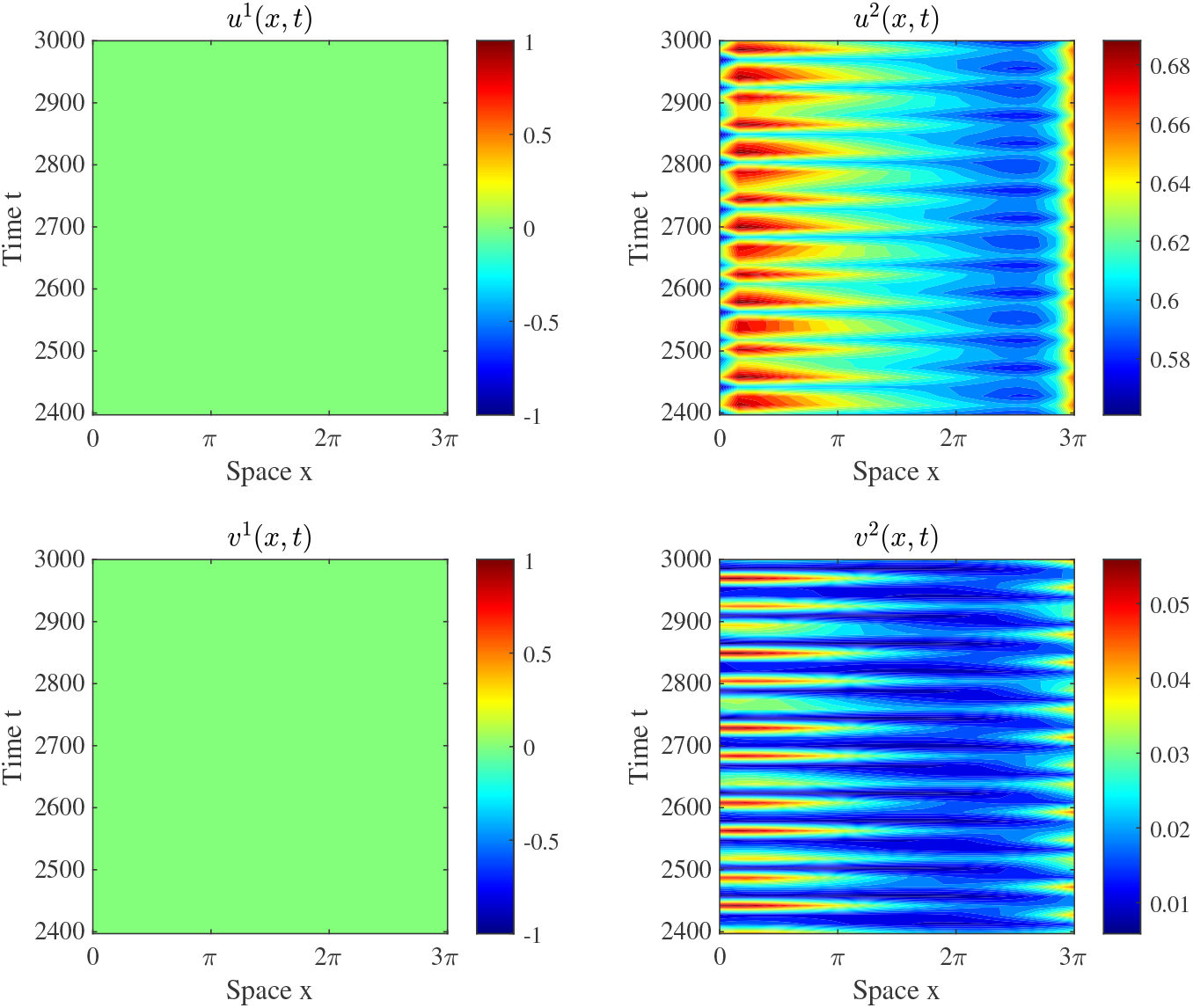
The solution of (1) with parameters given in (48) and (49).

The minimum value of *G* also occurs at ({2}, {2}), as computed that

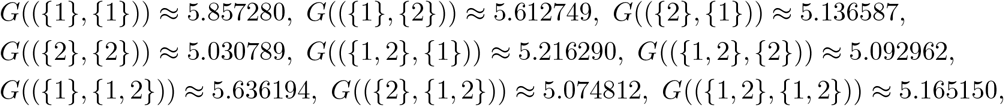

Thus, the combination closest to that at which *G* attains its minimum has competition advantages.

## 8 Discussion: The concept of IFD in a time-periodic environment

In temporally constant environments, a population is said to achieve an ideal free distribution (IFD) if the fitness (as a function of *x* only) is equilibrated. The evolutionary stability of IFD for a single population has been established across many modeling contexts, see Cantrell et al. (2017, 2012, 2022); Maciel et al. (2020); Cantrell et al. (2007) and references therein, where it is proved that when a single species population achieves IFD, then mutant phenotypes cannot invade when rare.

When considering a community of ecologically identical species, then the fitness of each resident species depends both on the abiotic environment and the other resident species it is interacting directly or indirectly with, and a community of species is said to achieve IFD if the fitness of every species is equilibrated in space. The evolutionary stability of IFD is proved in Cantrell et al. (2007) in the setting of discrete-patch models, where a community of interacting species achieving IFD is shown to resist invasion by rare mutant phenotypes. Hence, an ecological community forms an evolutionary endpoint if the community as a whole achieves IFD.

In this paper, we establish the evolutionary stability of IFD in the context of a system of reaction-diffusion equations modeling a community of multiple prey and predator species. Our analysis reveals the following biological insights:

### IFD in time-periodic environments

Local ecological conditions are not always temporally constant. For temporally periodic environments, fitness of a given population is generally a function of *x* and *t*, and it is demonstrated that the appropriate notion of IFD is for the fitness to equilibrate in *x*, but not necessarily in *t*. Inspired by our previous work in competition systems in the reaction-diffusion-advection model setting (Cantrell et al. 2021) and in patch model setting (Lam and Zhang 2025), we show here that an appropriate notion of IFD is introduced where we require that fitness be equilibrated in space alone, but not necessary in time (see Definitions 1 and 2). Using this definition as an ansatz, we determine analytically the IFD based on environmental parameters. These conditions depend on the positive solutions of a non-autonomous predator-prey ODE system, which reflects the fact that anticipation based on nonlocal information, is needed for a population to achieve IFD in an environment that is varying both spatially and temporally. Such nonlocal information can be obtained based on e.g. a trial-and-error manner or via memory-based dispersal (Shi et al. 2021). For instance, empirical evidence shows that some whales can accurately track the places and times where resources will be most reliably abundant over periods of years, using memory and resource tracking to effectively average out yearly variations (Abrahms et al. 2019). Interestingly, species can still over-match or under-match resources when considering the time-periodic environments. We also discuss the necessary and sufficient conditions for a joint IFD to be feasible (see Theorem 9). It turns out that there exist time-periodic environments in which IFD is impossible no matter what dispersal strategies are adopted. This is due to the presence of “generalized sinks” where the species is too much constrained by its birth and death rates and cannot be abundant enough to equilibrate fitness.

### IFDs can be evolutionary endpoints for communities

We show that joint IFD consisting of a large number of species can be evolutionarily stable (see Theorems 10 and 11, generalizing previous results to reaction-diffusion-advection context. This offers a plausible mechanism for the maintenance of food webs in nature consisting of a large number of interacting species. However, when two different coalitions of species can form IFD, we cannot determine which subset can dominate. We conjecture that, under more realistic stochastic conditions, the two subcommunities of species form neutral populations, which remain stable while, over time, individual subcommunities will drift according to random walk within this constraint of global IFD, and the number of such neutral groups will typically decrease over time until one neutral group remains (Chesson and Huntly 1997; Hubbell 2011).

### IFDs are neighborhood invader strategies

Going a step beyond local stability analysis performed in Cantrell et al. (2007), our argument shows the global stability of joint IFD formed in a community. Namely, if a subset of species can achieve IFD, then the whole community must approach an IFD, even if the initial data is a large perturbation of IFD. Roughly speaking, our results show that a new species can invade a community and establish a viable population if it complements the existing population so that, as a whole, the community including the new species can achieve IFD.

### Non-IFD is selected against

When the community does not form a joint IFD, then we show that it can be invaded by exotic species, provided it can employ the appropriate dispersal strategy (see Theorem 2). This shows that IFD is a necessary condition for a community to be evolutionarily stable.

### Selection advantage of approximate IFD

Extending and going beyond our analytical results, we also investigated the situation when IFD is not achieved. For this purpose, we introduced in Section 7 a measure of how “close” a community is to achieving an IFD. Our numerical results show that a new species is more likely to invade and establish a viable population within an existing community if its presence can bring the community closer to achieving IFD. Interpreted in the framework of adaptive dynamics, it follows that by repeated introduction of mutant phenotypes (either by immigration or by genetic diversification), nature selection tends to shape the community of species towards IFD. In the framework of adaptive dynamics, this suggests that the IFD is convergence stable as well as evolutionarily stable.

## Declarations

- **Funding** RSC and CC are supported partially by National Science Foundation (DMS 18-53478). KYL is supported partially by National Science Foundation (DMS-2325195); HZ is supported by National Natural Science Foundation of China (No. 12401213, 12571210) and China Postdoctoral Science Foundation (No. GZB20240437).
- **Conflict of interest** The authors declare that they have no conflict of interest.
- **Ethical Approval** This declaration is not applicable.
- **Data Availability** All data generated or analyzed during this study is included in this article.

## Appendix A

**The proof of Lemma 7**

*The proof of Lemma 7* Let *A*_*i*_ and *B*_*ij*_, *i* = 1, 2, *j* = 1, 2, be defined as (8). We first prove statement (i). Since *a, b, c, d, e, f >* 0 are independent of *t*, it follows from (8) that *A*_*i*_ and *B*_*ij*_ *>* 0 are constants. We then have

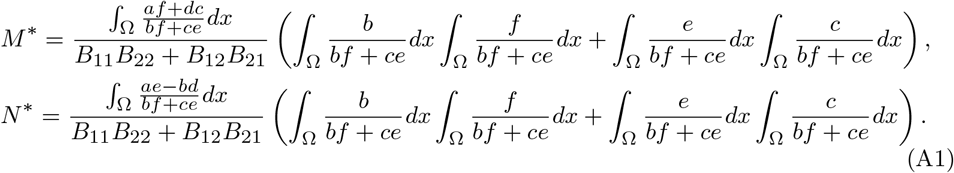

Therefore, *M* ^*^, *N* ^*^ are positive constants provided 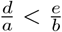.

To prove (ii), define 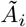 and 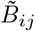 by replacing *a, b, c, d, e, f* with 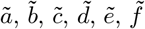 in *A*_*i*_ and *B*_*ij*_, respectively. Then (7) becomes a fast-slow time-periodic system:

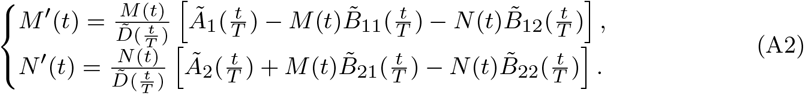

Set 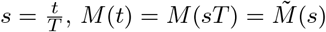 and 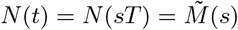. This yields

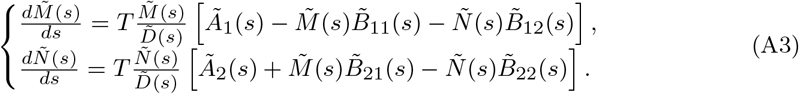

where 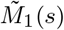 and 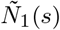 solve (A5). Substituting this into (A3) and comparing coefficients of like powers of 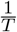, we obtain

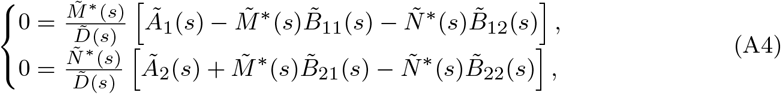

and

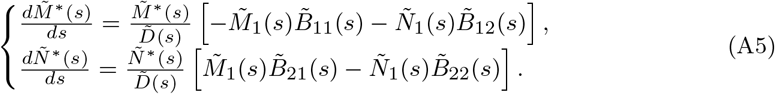

Clearly, if 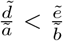, then (A4) has a unique 1-periodic solution 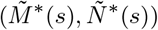 given by (A1) with *B*_*ij*_ and *a, b, c, d, e, f* replaced by 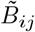 and 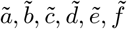, respectively. From (A5), 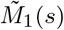 and 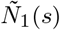 are uniformly bounded in *s*. Hence 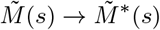 and 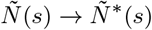 as *T* → ∞. Consequently, (7) has a positive *T*-periodic solution (*M*(*t*), *N*(*t*)) as *T* ≫ 1.

In the case of (iii), (*M*(*t*), *N*(*t*)) satisfies

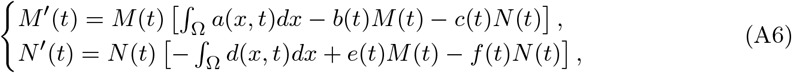

It suffices to prove that condition

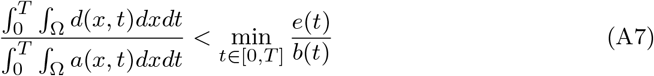

Implies 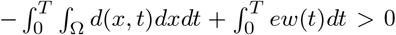, where *w*(*t*) is the unique positive *T*-periodic solution to *w* (*t*) = *w*(*t*) ∫_Ω_ *a*(*x, t*)*dx* − *b*(*t*)*w*(*t*). If this holds, the existence of a positive *T*-periodic solution (*M*(*t*), *N*(*t*)) follows directly from Corollary 3 (Ding et al. 1995).

Observe that 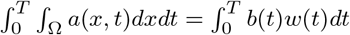. Then (A7) implies that

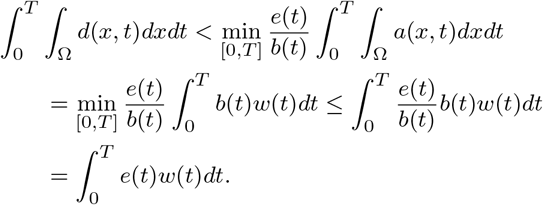

This completes the proof. □

## Footnotes

1 Note that for any compact set 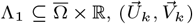 is well-defined on Λ_1_ for all *k* ≫ 1. Hence it makes sense to speak of precompactness of 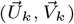 in 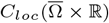.

